# Effect of assortative mating and sexual selection on polygenic barriers to gene flow

**DOI:** 10.1101/2024.07.30.605898

**Authors:** Parvathy Surendranadh, Himani Sachdeva

## Abstract

Assortative mating and sexual selection are widespread in nature and can play an important role in speciation, through the buildup and maintenance of reproductive isolation (RI). However, their contribution to genome-wide suppression of gene flow during RI is rarely quantified. Here, we consider a polygenic ‘magic’ trait that is divergently selected across two populations connected by migration, while also serving as the basis of assortative mating, thus generating sexual selection on one or both sexes. We obtain theoretical predictions for divergence at individual trait loci by assuming that the effect of all other loci on any locus can be encapsulated via an effective migration rate, which bears a simple relationship to measurable fitness components of migrants and various early generation hybrids. Our analysis clarifies how ‘tipping points’ (characterised by an abrupt collapse of adaptive divergence) arise, and when assortative mating can shift the critical level of migration beyond which divergence collapses. We quantify the relative contributions of viability and sexual selection to genome-wide barriers to gene flow and discuss how these depend on existing divergence levels. Our results suggest that effective migration rates provide a useful way of understanding genomic divergence, even in scenarios involving multiple, interacting mechanisms of RI.

## 1. Introduction

Reproductive isolation (RI) typically results from the interaction of multiple processes that cause loss of hybrid fitness, thus reducing gene flow between populations. Ecological specialisation arising from adaptation to different environmental niches, immigrant inviability, or intrinsic genetic incompatibilities may all generate postzygotic barriers to gene flow and maintain genetic differences between populations (Rundle and Nosil (2005); Coughlan and Matute (2020); Rice and Hostert (1993)). Additionally, processes such as assortative mating and sexual selection can act as both prezygotic and postzygotic barriers— by reducing heterospecific matings (that produce hybrids) as well as the mating success of hybrids. Indeed, it has been argued that the buildup of RI typically involves coupling between both postzygotic and prezygotic barriers (Butlin and Smadja (2018)).

Assortative mating (AM)– which involves positive phenotypic (and genetic) correlation between mating pairs (Lewontin et al. (1968)), is especially common in animals Jiang et al. (2013). Numerous studies have documented its role during RI, e.g., in cichlids (Elmer et al. (2009); Stelkens and Seehausen (2009)), swordtail fish (Schumer et al. (2017)), sticklebacks (Vines and Schluter (2006)), butterflies (Jiggins et al. (2001)), and marine snails (Johannesson et al. (2008)). AM decreases heterozygosity and genic variance within a population, but simultaneously increases total genetic variance (by generating higher levels of linkage disequilibria (LD)) (Lynch et al. (1998)), thus increasing the potential for divergence and speciation.

AM may occur either due to temporal or spatial separation of population groups, via preference-trait mechanisms, or via phenotype matching between potential mates (Kopp et al. (2018); Jiang et al. (2013); Kirkpatrick and Ravigné (2002)). Each mechanism differs in the extent of sexual selection it generates— ranging from strong sexual selection under phenotype matching to almost none when assortment is due to temporal or spatial separation. Sexual selection typically reduces mating success of rare phenotypes, and thus may counter the effects of AM (Servedio and Boughman (2017); Safran et al. (2013)). For example, when AM involves sexual selection on both sexes (e.g. if both males and females suffer reduced mating opportunities by being choosy), then its net effect is to generate stabilizing selection and inhibit divergence (Kirkpatrick and Nuismer (2004)).

Numerous studies have examined how different isolating mechanisms, namely assortative mating (Doebeli (1996); Kondrashov and Shpak (1998); Kopp et al. (2018)), ecological selection (Smith (1966); Maan and Seehausen (2011)), and sexual selection (Servedio (2016); Servedio and Kopp (2012); Schumer et al. (2017)) influence divergence and RI in the presence of ongoing gene flow. These suggest that RI is facilitated by coupling between traits that act as reproductive barriers (Coyne and Allen Orr (2004); Smadja and Butlin (2011); Kirkpatrick and Ravigné (2002)). Such coupling can arise if loci underlying different traits have pleiotropic effects or are tightly linked and not easily broken apart by recombination (Felsenstein (1981); Butlin and Smadja (2018); Smadja and Butlin (2011); Servedio (2009)). In this regard, speciation is thought to be most effective with magic traits, i.e., if the same trait is under divergent selection and also mediates AM. Magic traits are now known to be more common than previously thought (Servedio et al. (2011a)): well-known examples include wing color pattern in Heliconius (Merrill et al. (2012)) and body size in Littorina (Perini et al. (2020)).

Magic traits may differ in their mechanism of AM and the extent to which they are under sexual selection, and have been studied extensively to understand the evolution of mate choice during speciation (Servedio et al. (2011b); Servedio and Kopp (2012); Servedio and Boughman (2017)). While previous work has mostly focused on few-locus models of mate choice, here we consider a polygenic trait influenced by many small-effect alleles. Not only is this more realistic in view of the polygenic architecture of most traits but also much less understood, with previous studies relying largely on simulation of two populations (Sachdeva and Barton (2017); Muralidhar et al. (2022)) or of hybrid zones (Irwin (2020); Perini et al. (2020)). In general, the evolution of polygenic divergence and RI depends on the ease with which *multiple* locally advantageous alleles establish and/or are maintained across the genome despite maladaptive gene flow between populations. This in turn depends on whether selected loci evolve more-or-less independently or if LD between sets of maladapted alleles causes them to be eliminated *together* before they recombine onto fitter backgrounds. This was first studied in the context of hybrid zones, where it was found that the strength of LD between multiple selected loci depends on the ratio of total selection and total recombination (Barton (1983)).

As we demonstrate here, a powerful way of analysing the effects of LD on divergence is by tracking how introgressing alleles at any one trait locus are transferred between genetic backgrounds, typically via multiple recombination events over multiple generations. The different genetic backgrounds on which the focal allele finds itself may differ in their hybrid index (i.e., the proportion of genetic material they derive from the parental populations) and thus also have different fitness values, which influences the transmission probability of the allele. This is captured by the notion of the *effective* migration rate (*m*_*e*_), which is the rate at which neutral introgressing alleles are transferred between hybridising populations (Bengtsson (1985)). In essence, strong LD between large numbers of loci reduces *m*_*e*_ across the genome (Barton and Bengtsson (1986)). This increases divergence between hybridising populations, which reduces hybrid fitness, further decreasing effective migration rates. Indeed, it has been argued that this genomewide reduction in *m*_*e*_ between hybridising populations provides a natural way of quantifying RI (Westram et al. (2022)). Subsequent work has generalised theory based on *m*_*e*_ to weakly selected loci with partial and/or transient divergence between populations (Sachdeva (2022)), heterogeneous effect sizes and dominance (Zwaenepoel et al. (2024)) and sex-linkage (Fräisse and Sachdeva (2021)). However, all of this work is restricted to randomly mating populations.

In this paper, we extend theory based on effective migration rates to a scenario with assortative mating due to female preference for phenotypically similar males, which leads to sexual selection on one or both sexes. The phenotype underlying assortment is polygenic and under divergent selection across a mainland and island: thus, we have a simple ‘magic trait’ scenario of speciation. We derive expressions for *m*_*e*_ at an unlinked neutral locus and then use this to predict allele frequency divergence at individual trait loci under migration-selection-assortment balance by assuming that the effect of all other loci on the focal locus is encapsulated by this effective migration rate. This allows us to predict mean trait divergence between mainland and island, and explore how assortment, viability and sexual selection jointly influence the critical threshold at which migration swamps adaptive divergence. We then use this to disentangle the relative contributions of viability and sexual selection to the genomewide barrier to gene flow in parameter regimes where partial RI persists despite limited hybridisation.

## 2. Model and Methods

### 2.1 Model

Consider an island subject to one-way migration from a large mainland, such that a fraction *m* of individuals on the island are replaced by migrants in every generation. Individuals are haploid and hermaphroditic (i.e. produce both male and female gametes), and can thus play the male or female role during reproduction. Each individual expresses an additive polygenic trait 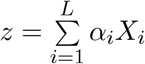 influenced by a large number *L* of unlinked, biallelic loci, where *X*_*i*_ = 0, 1 denote alternative alleles and *α*_*i*_ *>* 0 is the effect size at locus *i*. The trait is under directional selection on the island– the viability component of fitness of an individual with trait value *z* is proportional 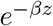,where *β* denotes the strength of viability selection. We will not explicitly model the mainland population but assume that trait values follow a normal distribution, or alternatively, that migrant genotypes are maximally deleterious on the island (see sec. 2.2). Unless stated otherwise, we will also assume that the island is initially perfectly adapted (with *z* = 0 for each individual).

Individuals on the island mate assortatively, with trait *z* also serving as the basis for AM via female choice. We assume a Gaussian choice function, such that the probability of mating between a male and female with trait values *z* and *z* is proportional to 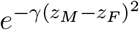,where *γ* is the strength of assortment. In other words, females preferentially mate with males with trait values within a few 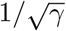 of their own. Thus, the assortment trait is a ‘magic trait’, which simultaneously influences mate choice and is under both natural and sexual selection, thereby generating both postzygotic and prezygotic barriers to gene flow.

We consider two different models of AM– allowing for sexual selection on both sexes (Model I) or only on males (Model II)– sometimes referred to as the ‘plant model’ and ‘animal model’ of assortment (Kirkpatrick and Nuismer (2004)). The probability *M* (*z*_*M*_, *z*_*F*_) of mating between a male and female with trait values *z*_*M*_ and *z*_*F*_ under the two models is:

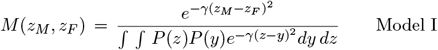

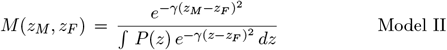

Here, *P* (*z*) is the trait value distribution just before mating. Note that under Model II, mating probabilities are normalized such that any female, regardless of trait value, mates with probability 1, while under Model I, females with rare phenotypes have lower mating probability. Thus, sexual selection acts only on males in the former case but on both sexes in the latter.

### 2.2 Effective migration rate at a neutral locus

Our analysis is based on effective migration rates (*m*_*e*_)– defined as the rate at which neutral alleles entering a population via migration are incorporated into the resident gene pool (Bengtsson (1985); Barton and Bengtsson (1986), where ‘residents’ are defined more precisely later. In our model, migrants and their descendants experience both viability and sexual selection and are less fit than residents. Consequently, in order to establish, neutral alleles must recombine away from migrant genetic backgrounds before these are eliminated by selection. Thus, *m*_*e*_ will vary along the genome, being lower at sites that are tightly linked to a selected locus or in genomic regions with a high density of selected loci. However, we will focus on *m*_*e*_ at a neutral locus that is *unlinked* to any selected locus, since we are interested in quantifying the average or genomewide reduction in gene flow.

Under rare migration (*m* ≪ 1), the ratio *m*_*e*_*/m* for an unlinked neutral allele, also called the gene flow factor *g*, is equal to the reproductive value (RV) of migrants relative to residents (Kobayashi et al. (2008)). Here, RV refers to the long-term genetic contribution of the migrant in the recipient population (Fisher (1999); Barton and Etheridge (2011)), and is given by:

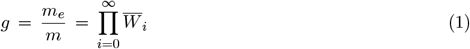

where 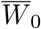 denotes the average fitness of migrants relative to residents, and 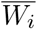 the average relative fitness of *i*^*th*^ generation descendants of migrants (see, e.g., Westram et al. (2022)). For *m* ≪ 1, individuals with recent immigrant ancestry are rare, so that migrants and their descendants mate primarily with residents. As a result, first-generation descendants of the migrants are *F*_1_ hybrids, second-generation descendants are typically first-generation backcrosses (with residents), *i*^*th*^ generation descendants are (*i*−1)^*th*^ generation backcrosses (denoted by *BC*_*i*−1_), and so on. Thus, computing *m*_*e*_ boils down to calculating the relative fitness of successive back-crosses.

In the following, we classify individuals on the island by the number of generations leading back to their most recent immigrant ancestor, i.e., into *F*_1_, *BC*_1_, *BC*_2_, up to *BC*_*n*_ (where *n* is arbitrary). All other individuals, i.e., those with no migrant ancestor in the previous *n* + 1 generations, are designated as ‘residents’. We further assume trait values to be normally distributed *within* any group of individuals with the same level of recent migrant ancestry. In other words, the trait distributions *P*_*r*_(*z*), *P*_1_(*z*), … *P*_*i*_(*z*), … amongst residents, *F*_1_ hybrids, *i*^*th*^ generation descendants (who are *BC*_*i*−1_) and so on, are assumed to be normal with means 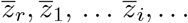 and variances *V*_*r*_, *V*_1_, … *V*_*i*_, … respectively (see figure 1A; also Table 1). For generality, we also take the migrant trait value distribution to be normal with mean 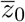 and variance *V*_0_. However, in the Results, we will only consider a scenario where the mainland is fixed for alleles locally deleterious on the island (so that 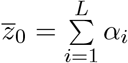 and *V*_0_ = 0).

**Table 1:** Key parameters and variables.

| Parameter/<br>Variable | Description |
| --- | --- |
| $z$ | Additive magic trait; individual trait value $z = \sum_{i=1}^L \alpha_i X_i$,<br>where $X_i = 0$ or $1$ denote alternate alleles at locus $i$ |
| $\alpha_i$ | Effect size at a locus $i$ |
| $L$ | Number of loci influencing magic trait |
| $N$ | Population size |
| $m$ | Rate of migration from mainland to island |
| $\beta$ | Strength of viability selection on island; individual with trait value $z$ has viability $\propto e^{-\beta z}$ |
| $\gamma$ | Strength of assortative mating |
| $M(z_M, z_F)$ | Probability of mating between a male and female with trait values $z_M$ and $z_F$;<br>differs under the two models of assortative mating |
| $s_i$ | Selection coefficient at locus $i$ given by $s_i = \beta \alpha_i$; we consider equal-effect loci with $s_i = s$ |
| $m_e$ | Effective migration rate at any (equal-effect) trait locus |
| $g$ | Gene flow factor $g = m_e/m$ |
| $\Delta$ | Mean trait divergence between mainland and island, $\Delta = \sum_{i=1}^L \alpha_i(1 - p_i)$; $p_i$ is frequency of<br>locally deleterious allele at locus $i$ on island; mainland is fixed for the alternative allele |
| $\Delta_0$ | Maximum possible divergence between mainland and island, given by $\alpha L$ |
| $Y$ | Allele frequency divergence per locus, $Y = \Delta/\Delta_0$, equals $1 - p$ for equal-effect loci |
| $\beta \Delta_0, \gamma \Delta_0^2$ | Scaled strength of viability and sexual selection; amount by which (log) fitness of<br>migrants is reduced due to viability and sexual selection respectively, when $\Delta = \Delta_0$ |
| $m_c/s$ | Critical migration rate (relative to selection per locus) at which divergence $Y$ becomes 0 |
| $Y_c$ | Allele frequency divergence associated with the tipping point |
| **Quantities that appear in derivation of gene flow factor $g$** |  |
| $P_r(z), P_i(z)$ | Trait value distribution among residents and $i^{th}$ generation descendants of migrants;<br>$i = 0$ corresponds to migrants. |
| $\bar{z}_r, \bar{z}_i$ | Mean trait value among residents and $i^{th}$ generation descendants of migrants |
| $V_r, V_i$ | Variance of trait values among residents, $i^{th}$ generation descendant of migrants |
| $V_{r,r}, V_{r,i}$ | Within-family variance when one parent is a resident and the other a resident<br>or an $i^{th}$ generation descendant of a migrant |
| $\bar{W}_i$ | Mean fitness of $i^{th}$ generation descendants of migrants relative to residents |

**Figure 1:**
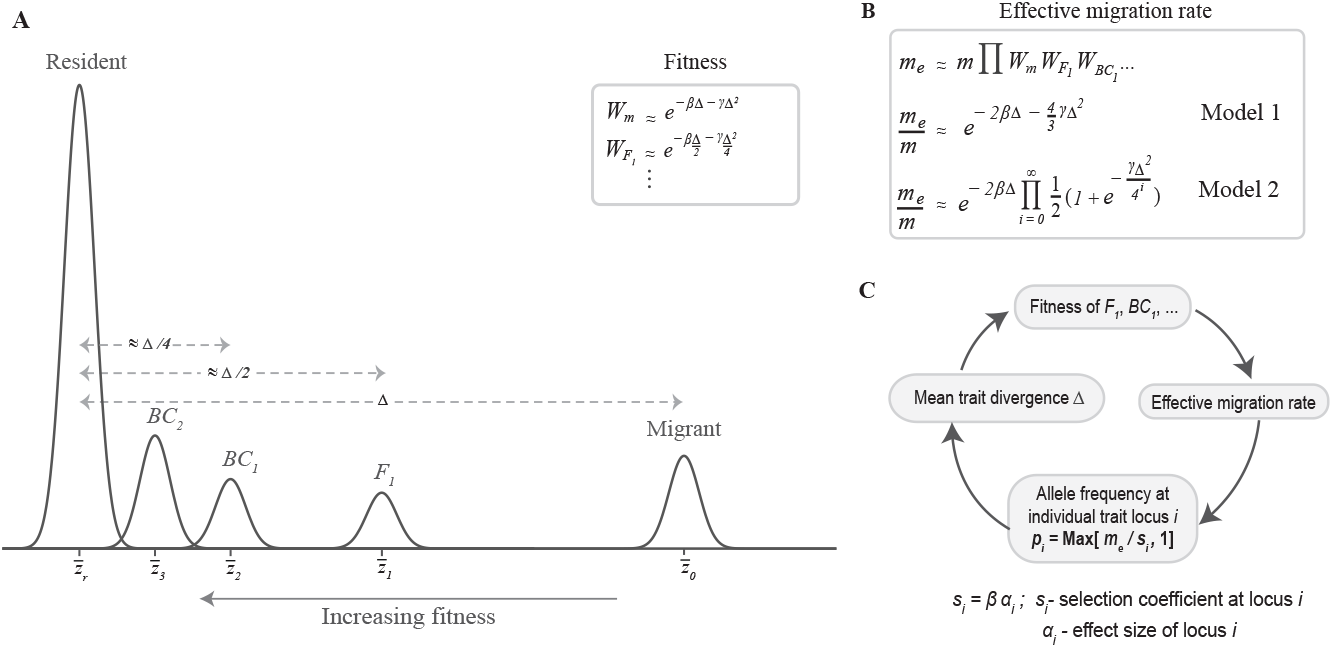
Schematic summarising our theoretical approach for predicting mean trait divergence Δ (or allele frequency divergence per locus *Y*) using effective migration rates (*m*_*e*_). **A**: Distribution of trait values on the island is the sum of trait value distributions associated with residents, *F*_1_s, *BC*_1_s, and so on, where each such distribution is assumed to be approximately normal. Denoting the difference between trait means of migrants and residents by 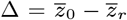, the difference between trait means of *F*_1_s and residents is ≈ Δ*/*2, and more generally, between that of *i*^*th*^ generation descendants (of migrants) and residents is Δ*/*2^*i*^. This allows us to calculate *average relative fitness* of *F*_1_s, *BC*_1_ etc. (inset). **B**: The *gene flow factor g* (which describes the reduction in neutral gene flow at a site unlinked to any selected site) can be calculated from the average relative fitness of migrants and of individuals with recent migrant ancestry (*F*_1_s, *BC*_1_s), as in eq. 1. This gives approximate expressions for *g* under Model I (sexual selection on both sexes) and Model II (sexual selection only on males) **C**: The *mean trait divergence* Δ can be calculated by first using *m*_*e*_ to predict the allele frequency *p*_*i*_ at trait locus *i* = 1, …, *L*, then summing over loci to obtain the mean trait divergence Δ, then using this to predict relative fitnesses of *F*_1_s, *BC*_1_s and so on (as in A.), and finally using these to predict the effective migration rate *m*_*e*_ (as in B.). This kind of circular dependence, where *m*_*e*_ depends on Δ which depends on *m*_*e*_ (via allele frequencies), allows for a self-consistent solution for the mean trait divergence Δ.

Assuming normally distributed trait values is justified for traits influenced by a large number of loci of small effect. The inheritance of such traits is described by the infinitesimal model (Fisher (1918)), which states that trait values of the offspring of any two individuals are normally distributed about the mean of the parents, with a ‘within-family’ variance *V*_∗_, which depends only on the genetic relatedness between parents, regardless of selection, non-random mating etc. (Barton et al. (2017)). We further assume that *V*_∗_ depends only on the extent of migrant ancestry of the parental individuals, and does not vary significantly across (for instance) different resident *× F*_1_ parental pairs, all of which thus have approximately the same *V*_∗_. We denote the within-family variance by *V*_*r,r*_ when both parents are residents, *V*_*r*,0_ when one parent is a resident and the other migrant, and *V*_*r,i*_ when one parent is a resident and the other an *i*^*th*^ generation descendant of a migrant.

Then, we can express the distribution *P*_*i*_(*z*) in terms of the parental trait value distributions *P*_*r*_(*z*_1_) and *P*_*i*−1_(*z*_2_) (eq. 2a). Further, we can express the mean fitness 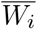 of *i*^*th*^ generation descendants (relative to residents) as an integral over the parental distributions (eq. 2b).

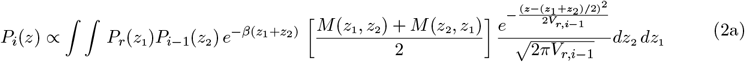

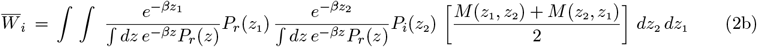

In eq. 2a, *z*_1_ and *z*_2_ denote respectively the trait values of the resident parent and the parent who is an (*i* − 1)^*th*^ generation descendant of descendant of a migrant, with corresponding trait distributions *p*_*r*_ (*z*_1_) and *p*_*i* −1_ (*z*_2_). The term 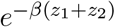 captures the effect of viability selection on the parents, while *M* (*z*_1_, *z*_2_) (which is the probability of mating between a male and female with trait values *z*_1_ and *z*_2_ respectively) captures the effect of AM and sexual selection. Note that the non-resident parent may have a different probability of being chosen as a mate depending on whether it takes the female or male role during reproduction: this is captured by the two terms *M* (*z*_1_, *z*_2_) and *M* (*z*_2_, *z*_1_) (that are unequal under Model II). Finally, the last term gives the distribution of trait values of the offspring under the infinitesimal model: it captures the effect of recombination between parental genotypes and random segregation. Supplementary Information (SI)A provides a detailed derivation of eq. 2.

Integrating over eq. 2a gives an expression for the mean and variance of trait values among the *i*^*th*^ generation descendants in terms of the means and variances among (*i*−1)^*th*^ generation descendants and among residents. This specifies a set of recursions for 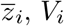 and *V*_*r,i*_ which can be solved numerically (see SI, **A**). In turn, these predict the relative fitnesses 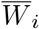 (using eq. 2b), from which one can calculate the gene flow factor *g* (using eq. 1) and *m*_*e*_.

However, in the main paper, we will derive a more approximate and intuitive expression for *g* by simply tracking the difference between trait means of residents and other groups (i.e., *F*_1_s, *BC*_1_s and so on), while neglecting phenotypic variance *within* any group. More concretely, we assume that the mean trait value of hybrid offspring is the average of the trait means of the parental subgroups. Thus, for example, *F*_1_ hybrids are assumed to be midway between residents and migrants, giving 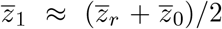,which gives: 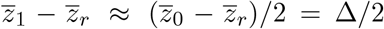.Here 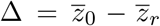 denotes the difference in the trait means of migrants and residents, which we refer to as mean trait divergence. It follows similarly that 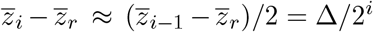 (see figure 1A). Thus, in effect, every successive generation of backcrossing halves the average phenotypic distance of the (next-generation) backcross from the resident phenotype (Sachdeva (2022); Zwaenepoel et al. (2024)).

As we argue below and in SI, **A**, this simple argument holds as long as *βV*_*i*_ ≪ Δ and *γV*_*i*_ ≪ 1 (for all *i*). Moreover, under these conditions, the relative fitness of any group of descendants is (from eq. 2b): 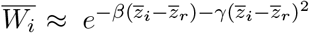 for Model I (sexual selection on both sexes) and 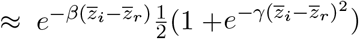 for Model II (sexual selection only on males). Now, using 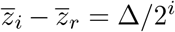, where Δ is the mean trait divergence between mainland and island, we obtain the following approximate expressions (see figure 1B) for the gene flow factor 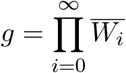:

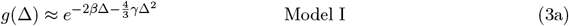

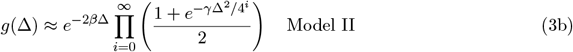

A more detailed derivation of eq. 3 is provided in SI, **A**. A striking feature of eq. 3 is that under both models of sexual selection, the gene flow factor can be expressed as a product of two terms– the first describing the reduction in reproductive value of migrants due to viability selection, and the second due to sexual selection. The first term above, *e*^−2*β*Δ^, is the square of the ecological fitness of migrants relative to residents. The second term is related to the relative mating success of migrants and differs between the two models. For instance, under Model I, the sexual selection term is simply the relative mating success of migrants raised to the power 4*/*3. These powers (of 2 or 4*/*3) emerge from the specific fitness and mating functions assumed in our model. However, more generally, the fact that the viability and sexual selection components of *g* bear a simple relationship to the corresponding fitness components (of say migrants or *F*_1_s) is due to there being a simple relationship between the immediate fitness and long-term reproductive value of individuals when fitness is polygenic.

What assumptions underlie our simple approximation for trait means, which is the basis of eq. 3? First, within any group of descendants, say *F*_1_s, individuals closer to the resident phenotype will have higher viability, causing the trait mean of *BC*_1_s (the hybrid offspring of *F*_1_s and residents) to be slightly shifted towards residents rather than exactly midway between the two parental groups. The magnitude of this shift is proportional to *βV*_1_; thus, in ignoring the effects of viability selection *within* subgroups, we assume that *βV*_*i*_ ≪ Δ, where Δ is the mean trait divergence between mainland and island. As we argue in SI, **A**, under migration-selection balance, the various *βV*_*i*_ must be comparable to the migration rate *m*, and can thus be neglected if *m* ≪ 1.

Second, we ignore the effects of sexual selection within subgroups, which may lead to small differences in mating success among (say) *F*_1_s, again slightly shifting the *BC*_1_ trait mean towards one of the parental subgroups. Moreover, AM can also increase the phenotypic variance within each group (Fisher (1918); Wright (1921); Crow and Felsenstein (1968)), at least if sexual selection is weak, as under Model II (Kirkpatrick and Nuismer (2004)), further influencing the magnitude of these shifts.

As we argue in SI, **A**, both these effects can be ignored if *γV*_*i*_ ≪ 1, i.e., if the preference function is much wider than the typical phenotypic spread within any subgroup. Thus, in summary, our simple approximation for trait means 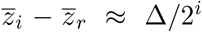 holds as long as *βV*_*i*_ ≪ Δ and if *γV*_*i*_ ≪ 1. In effect, these two conditions amount to assuming that viability and sexual selection are strong enough to affect the survival and mating success of migrants and their *F*_1_ and *BC* descendants relative to residents, but not strong enough to cause differential mating success *within* any group of descendants.

In summary, eq. 3 relates the gene flow factor *g* = *m*_*e*_*/m* at an unlinked locus to the mean trait divergence Δ between mainland and island at equilibrium, where Δ now depends on the allele frequency differences across all trait loci. In the following, we relate the allele frequency difference at any locus back to the effective migration rate *m*_*e*_, thus allowing us to obtain the mean trait divergence in a self-consistent way (see figure 1C).

### 2.3 Predicting allele frequency divergence at trait loci using *m*_*e*_

We now consider allele frequency dynamics at individual trait loci, assuming the mainland to be fixed for alleles that are deleterious on the island. First, consider the equilibrium allele frequency at a locus with effect size *α*_*i*_ under *linkage equilibrium*, i.e., neglecting the effects of LD between the focal locus and other trait loci. In SI, **B**, we show that sexual selection does not contribute to *direct* selection at a locus (at least under our assumed quadratic preference function), provided *α*_*i*_ is sufficiently small. More concretely, at migration-selection balance, the deleterious allele frequency *p*_*i*_ is simply *m/s*_*i*_, where *s*_*i*_ = *βα*_*i*_, provided *βα*_*i*_ ≪ 1 and *α*_*i*_ ≪ *γ/β* (SI, **B**). The latter condition can be re-written as *α*_*i*_*/*Δ ≪ (*γ*Δ^2^)*/*(*β*Δ)— this simply means that the relative contribution of any locus to trait divergence should be much smaller than the ratio of the sexual selection component of fitness to the viability selection component.

Following Sachdeva (2022), we now assume that the main effect of LD between immigrant alleles is to reduce *m*_*e*_ at any locus. This can be justified as long as trait loci are unlinked or weakly linked so that LD between these breaks down much faster than allele frequencies change. Further, if individual loci are weakly selected, then the effective migration rate at a trait locus is approximately equal to that at a neutral locus, so that in a large population, we have: 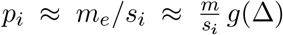. Here, the gene flow factor *g* is given by eq. 3, and captures the effect of both viability and sexual selection on migrants and their descendants.

Note that *g* depends on allele frequencies {*p*_*i*_, *i* = 1, ‥ *L*} at all *L* loci via the mean trait divergence 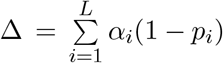. We can thus determine allele frequencies by numerically solving the *L* equations *p*_*i*_ = (*m/s*_*i*_)*g*(Δ), all coupled via Δ (Zwaenepoel et al. (2024)). However, here we will focus on the simpler case of equal-effect loci, such that *α*_*i*_ = *α* (and *s*_*i*_ = *s*) for all *i*. Then, in the absence of drift, equilibrium allele frequencies are equal across all loci (i.e., *p*_*i*_ = *p*), at least in parameter regimes where there is only one stable equilibrium. Thus, the mean trait divergence can be written as Δ = *αL*(1−*p*). In the following, we will express our results in terms of *Y* = Δ*/*Δ_0_, the trait divergence between the mainland and the island relative to the maximum possible divergence Δ_0_ = *αL*. Note that for the case of equal-effect loci, we have: *Y* = 1 − *p*, so that *Y* is also the mean allele frequency divergence per locus. It follows from eq. 3 that *Y* satisfies:

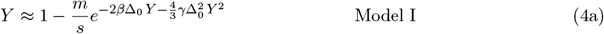

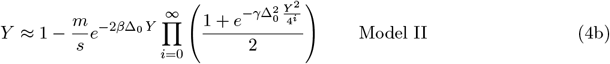

as long as a solution with 0 ≤ *Y* ≤ 1 exists, and is zero otherwise. Eq. 4 can be solved numerically to obtain allele frequency divergence per locus *Y* or the trait divergence Δ. In the case of Model II, the product in eq. 4b converges rapidly and can be computed by taking ∼ 20 terms.

Equation 4 is deterministic, implicitly assuming that *Ns* ≫ 1, where *N* is the size of the island population. However, our basic approach can be generalised to account for genetic drift, by treating each frequency as a (continuous) *random* variable described by a probability density *ψ*[*p*] (Sachdeva, 2022). Since a finite population subject to one-way migration requires non-zero mutation rates at trait loci in order to maintain long-term adaptive divergence, we allow for mutation at rate *µ* per individual per generation between alternative alleles at each locus. Then, under our approximation, the (unnormalised) equilibrium probability density *ψ*[*p*] is given by:

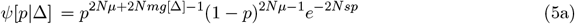

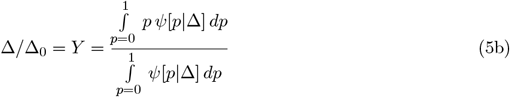

Equation 5a is simply the allele frequency distribution at a single locus at migration-selection-mutation-drift equilibrium, as predicted by the diffusion approximation (Wright (1937)), but with *m* replaced by *m*_*e*_ = *m g*[Δ]. As before, the effective migration rate *m*_*e*_ (or equivalently the gene flow factor *g*) is a function of the trait divergence Δ and is given by eq. 3. We can obtain Δ, in turn, by summing over allele frequencies at all trait loci. With equal-effect loci, we have: Δ = Δ_0_*Y*, where *Y*, the expected allele frequency divergence per locus, is obtained by integrating over *ψ*[*p*] (eq. 5b), thus allowing us to self-consistently solve for Δ (as in the deterministic case).

### 2.4 Deterministic simulations

In the main paper, we analyse the deterministic limit and test our approximations (eqs. 3 and 4) against deterministic simulations based on the hypergeometric model, which describes the inheritance of a quantitative trait influenced by equal-effect loci (Barton (1992); Doebeli (1996)). In each generation, migration is followed by viability selection, which is followed by assortative mating (involving sexual selection on one or both sexes), and production of offspring via free recombination between parental genotypes. The mainland is fixed for alleles that are locally deleterious on the island. Additionally, unless stated otherwise, simulations are initialised with the island fixed for the locally adaptive allele at each trait locus.

Note that with equal-effect loci, the trait value *z* of any individual is just *αj*, where *j* is the number of locally deleterious alleles it carries. SI, **C** specifies the recursions that describe how the distribution *P* (*j*) (or alternatively, *P* (*z*)) changes in any generation under the combined effects of migration, (viability and sexual) selection, AM and random segregation. Recursions for *P* (*j*) are iterated until equilibrium, and allele frequency divergence per locus *Y* obtained using: 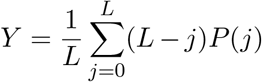. This can then be compared with theoretical predictions (eq. 4).

Individual-based simulations of finite populations (used to explore the effects of genetic drift on divergence) are described in SI, **D**.

## 3 Results

We now use predictions based on effective migration rates to investigate the combined effects of migration, viability and sexual selection, and assortative mating on the equilibrium allele frequency divergence between mainland and island as well as on the genome-wide reduction in neutral gene flow (as measured by the gene flow factor). In the main paper, we focus on the deterministic limit of the model; the effects of genetic drift are examined in SI, **D**. We first quantify how mean trait divergence declines with increasing migration (which is parameterized by *m/s*, migration rate relative to selective effect per locus), for different strengths of viability and sexual selection against migrants. This allows us to identify a critical migration threshold *m*_*c*_ beyond which local adaptation is swamped on the island, i.e., *Y* becomes zero. Next, we ask: how strong do viability and sexual selection have to be to shift this swamping threshold to *above* that for a single divergently selected locus? Finally, we examine the relative contributions of viability and sexual selection to RI under different parameter regimes, by using the increase in *F*_*ST*_ at an arbitrary site (unlinked to any trait locus) as a proxy for the genome-wide barrier effect.

Throughout, we quantify the strength of viability and sexual selection in terms of *β*Δ_0_ and 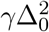,which denote respectively the reduction in log fitness of migrants (relative to residents) due to viability and sexual selection under conditions of maximum divergence between the mainland and island, i.e., when Δ = Δ_0_. If traits are highly polygenic (large *L* and small effect sizes *α*), then in the deterministic limit, divergence (as measured by *Y* = Δ*/*Δ_0_) depends essentially on the composite parameters *m/s, β*Δ_0_ and 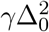,where Δ_0_ = *αL* (see eq. 4). However, for smaller *L* (say tens of loci), divergence levels may be sensitive to the exact genetic basis of the trait, i.e., depend on *α* and *L* separately (SI, **E**). In the main text, we only focus on the large *L* limit (where eq. 4 applies). Additionally, we assume a scenario of secondary contact where mainland and island are initially maximally diverged; other initial conditions (with partial divergence) are explored in SI, **F**.

### 3.1 Effect of viability and sexual selection on adaptive divergence

Figures 2A-2F show *Y*, the allele frequency divergence per locus (or equivalently, trait divergence between mainland and island relative to the maximum possible divergence) as a function of *m/s*, under the two models of AM in the deterministic limit. For each model, the plots depict divergence levels for *β*Δ_0_ = 0.2, 0.5, 1 (i.e., stronger viability selection from left to right). Further, for each *β*Δ_0_, we consider random mating (grey) as well as populations with increasing assortment (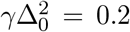 and 1), which generate increasing levels of sexual selection on both sexes (Model I; top) or only on males (Model II; bottom). The analytical predictions obtained by solving eq. 4 (lines) match well with the results of deterministic simulations of a trait influenced by *L* = 80 loci (points) unless net selection (viability and/or sexual selection) is very strong; see, e.g., red curves corresponding to 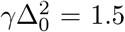 in figures 2E and 2F (with *β*Δ_0_ = 1). This discrepancy between simulations and theory becomes weaker with increasing number of trait loci (SI, **E**).

**Figure 2:**
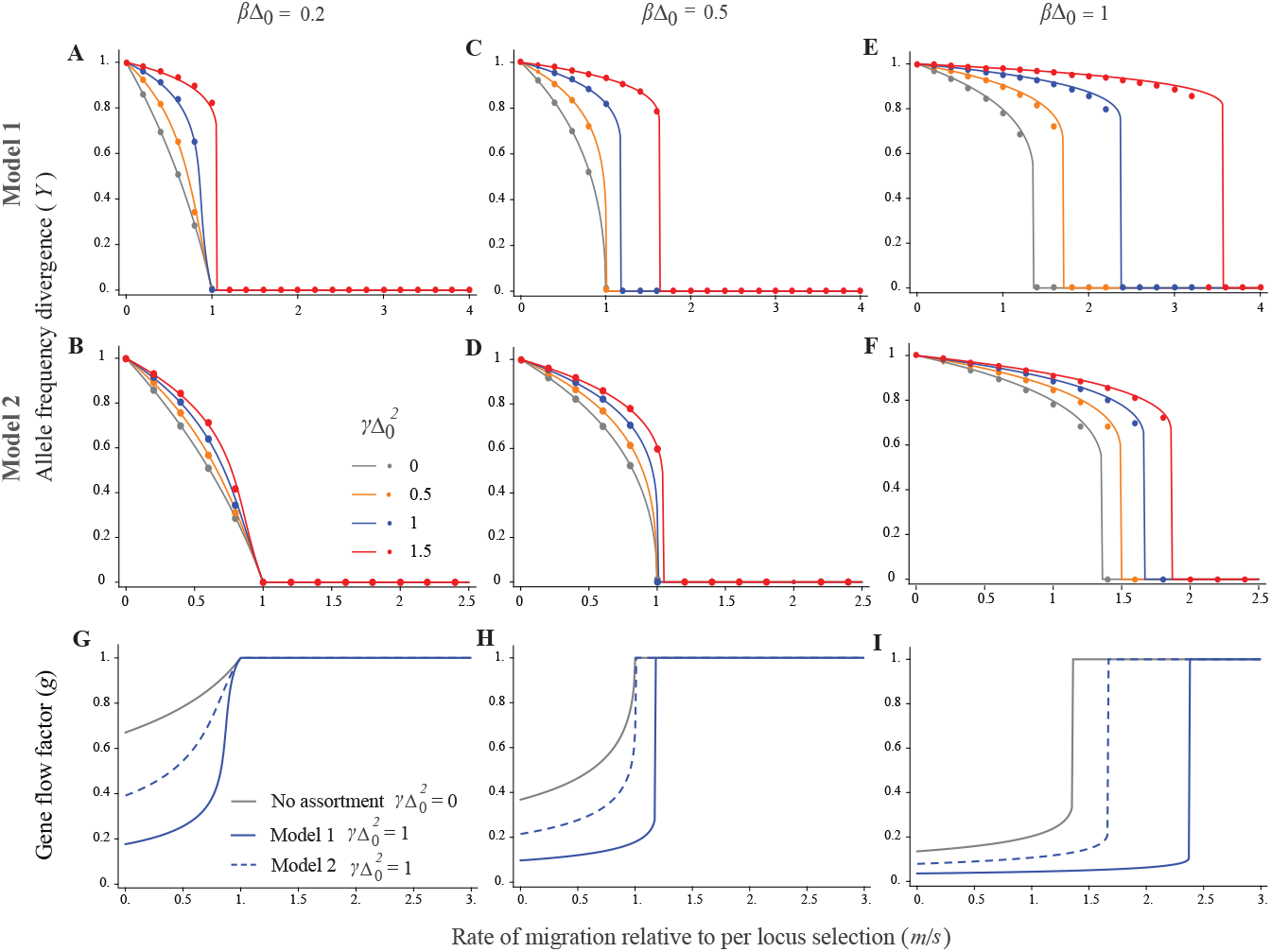
Allele frequency divergence per locus *Y* and gene flow factor *g* as a function of *m/s*, the migration rate relative to selection per locus. The top and middle rows depict respectively divergence levels for Model I (sexual selection on both sexes) and Model II (sexual selection only on males). Different subfigures within each row correspond to different strengths of viability selection (*β*Δ_0_ = 0.2, 0.5, 1 from left to right); different colors within each plot depict different strengths of AM and sexual selection (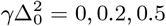,1 shown in grey, orange, blue and red respectively). Solid lines show theoretical predictions (obtained by numerically solving eq. 4) while dots represent the results of deterministic simulations of the hypergeometric model with *L* = 80 loci underlying the polygenic trait. Increasing viability and sexual selection cause sharper thresholds (‘tipping points’) for loss of divergence as well as an increase in the critical migration rate at which divergence is lost. The bottom row depicts theoretical predictions for the gene flow factor *g* for random mating (grey) vs. assortatively mating populations under Models I and Models II (solid vs. dashed lines), assuming 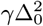.The gene flow factor and *m*_*e*_ are more strongly reduced under Model I than under Model II.

Thus, there is a significant range of parameter space where simple approximations based on *m*_*e*_ (eq. 4) predict equilibrium divergence accurately. Note that these approximations only account for the lower reproductive value of migrants due to sexual (and viability) selection, but ignore other effects of AM such as increased variance among residents, *F*_1_s, etc, and the increased probability of mating between individuals with recent migrant ancestry. That there is, nonetheless, close agreement between theory and simulations (which make no assumptions about AM), suggests that AM strengthens divergence in our model primarily by generating additional (sexual) selection against migrants and their descendants. This reduces *m*_*e*_, thus increasing the genomewide barrier effect and facilitating divergence (see also *Discussion*).

As evident from figures 2G-2I, this reduction in *m*_*e*_ (or equivalently *g*) is stronger under Model I (where sexual selection acts on both sexes) as compared to Model II (sexual selection only on males), and also depends on existing levels of divergence (which in turn depend on the strength of migration and viability selection).

Figure 2 also highlights how migration affects divergence differently in parameter regimes characterised by strong vs. weak selection against migrants. When sexual selection (due to assortment) and viability selection are weak, divergence declines *smoothly* with increasing *m/s*, going to zero at a critical threshold *m*_*c*_*/s* = 1, regardless of the actual value of *β*Δ_0_ and 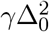 (e.g., in figure 2B). Note that *m*_*c*_*/s* = 1 is also the critical migration threshold for loss of adaptation at a single divergently selected locus. Thus, this threshold remains unchanged even if divergence is polygenic, as long as viability and sexual selection on the trait are weak.

By contrast, with strong viability and/or sexual selection, divergence at first decreases mildly with increasing migration, but then exhibits an *abrupt* collapse once it crosses a threshold *Y*_*c*_ (at a migration level *m/s* ≳ 1). Such a ‘tipping point’, defined as a discontinuous change in divergence (though not necessarily to *Y* = 0), can be observed in figure 2C and 2D at higher assortment and in 2E and 2F at all levels of AM. We can use eq. 3 to derive expressions for *Y*_*c*_, the allele frequency divergence per locus associated with tipping points under the two models of AM (details in SI, **G**):

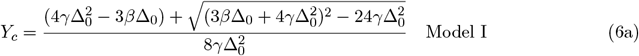

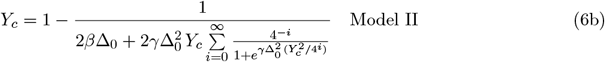

For Model II, eq. 6b specifies a transcendental equation for *Y*_*c*_, which can be solved numerically (e.g., by approximating the sum by the first ∼ 20 terms). Further, in SI, **G**, we show that the migration threshold associated with such a tipping point is: *m/s* = (1 − *Y*_*c*_)*/g*(Δ_0_*Y*_*c*_), where *g*, the gene flow factor, is given by eq. 3, and Δ_0_*Y*_*c*_ is the mean trait divergence at the tipping point.

The threshold *Y*_*c*_ can also be thought of as the minimum allele frequency difference per locus required to ‘lock in’ divergently selected alleles across the genome into distinct combinations that can be stably maintained by selection despite gene flow. Divergence levels lower than *Y*_*c*_ would result in an insufficient genomewide barrier to gene flow, causing a further decrease in divergence, further increasing *m*_*e*_ and so on, thus generating a tipping point. Interestingly, even though the tipping point occurs a specific level of migration (see, e.g., figure 2A-F), the divergence threshold *Y*_*c*_ associated with the tipping point is *independent* of this migration rate, and depends only on the strengths of viability and sexual selection (eq. 6).

In addition to generating tipping points, strong viability or sexual selection (or both) can also increase the critical migration threshold for loss of divergence *above* the single-locus threshold *m*_*c*_*/s* = 1 (see e.g., figures 2E, 2F). In other words, if selection against migrants and their descendants is sufficiently strong, then polygenic divergence can be maintained at migration levels at which a single divergently selected locus (in the absence of LD) would have been swamped.

Both tipping points and increased swamping thresholds may be explained as follows: higher levels of migration reduce adaptive divergence between mainland and island, thereby increasing *m*_*e*_ (see eq. 3) and weakening multi-locus selection against locally deleterious alleles, which increases their frequency– further reducing adaptive divergence and increasing *m*_*e*_, which finally results in the swamping of locally adaptive alleles above a certain migration threshold. Crucially, the positive feedback between loss of divergence at individual loci and increase in *m*_*e*_ across the genome is stronger when net selection against migrant phenotypes (either viability or sexual selection) is higher. In the next section, we will explore how strong do viability and sexual selection have to be to produce these effects?

Note that while figure 2 and the analyses below pertain to the deterministic model, the basic results also hold for a finite population of size *N*, provided *Ns* is large. For instance, tipping points occur under both models of AM if *Ns* ≳ 5, for moderately strong (viability plus sexual) selection (SI, **D**). Moreover, theoretical approximations based on *m*_*e*_ (eq. 5) furnish reasonably accurate predictions for divergence levels in finite populations as well (Figure S1, SI).

### 3.2 When does assortative mating and sexual selection lead to tipping points and higher swamping thresholds?

We first illustrate the various thresholds described above using a concrete example assuming weak viability selection (*β*Δ_0_ = 0.2) and sexual selection on both sexes (Model I). Under *weak* AM and sexual selection (in this example,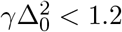), divergence decreases smoothly with increasing migration and is completely swamped at *m*_*c*_*/s* = 1: this behaviour is qualitatively similar to that of a single divergently selected locus. For *intermediate* assortment (here,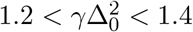), increasing migration leads to a tipping point or discontinuous change in divergence: this occurs once allele frequency divergence per locus approaches a threshold *Y*_*c*_ (given by eq. 6) and involves a strong reduction in the genomewide barrier effect. However, in this intermediate assortment regime, the tipping point still occurs at a migration rate lower than the single-locus swamping threshold *m*_*c*_*/s* = 1. For instance, for 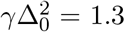,the tipping point occurs at *m/s* ≈ 0.95 and is associated with *Y*_*c*_ ∼ 0.6: thus, a tiny increase in *m/s* (say from 0.949 to 0.95) causes allele frequency divergence per locus to fall from 0.6 to 0.1 (Figure S5). A further increase in migration further reduces divergence, and complete swamping occurs (as before) at *m*_*c*_*/s* = 1. However, with even *stronger* assortment (in this example, 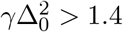), the genomewide barrier effect becomes strong enough to maintain divergence beyond *m/s* = 1, so that the tipping point now occurs at *m*_*c*_*/s >* 1. Further, divergence is completely swamped (*Y* falls to zero) at the tipping point. Thus, in this strong assortment regime, tipping points and swamping thresholds coincide and occur at a critical migration threshold that exceeds the single-locus threshold (Figure S5). This is in contrast to the intermediate assortment regime– where the tipping point occurs (i.e., divergence collapses to a low but non-zero level) at a migration rate which is lower than the swamping threshold.

We now ask: in what parameter regimes do we observe these qualitatively different behaviours? Figures 3A and 3B show the divergence threshold *Y*_*c*_ associated with tipping points (i.e., sharp drops in divergence), while figures 3C and 3D show the critical migration threshold *m*_*c*_*/s* beyond which divergence is completely swamped, as a function of the strength of viability selection (measured as *β*Δ_0_). As discussed above, the migration threshold associated with the tipping point is just the swamping threshold *m*_*c*_*/s* if assortment is strong. Figures 3A-3D show *Y*_*c*_ and *m*_*c*_*/s* for the two models of AM; colors in each plot correspond to different 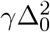,which quantifies the strength of sexual selection (due to AM). These predictions all follow from eqs. 3-6 (see SI, **G** for details).

**Figure 3:**
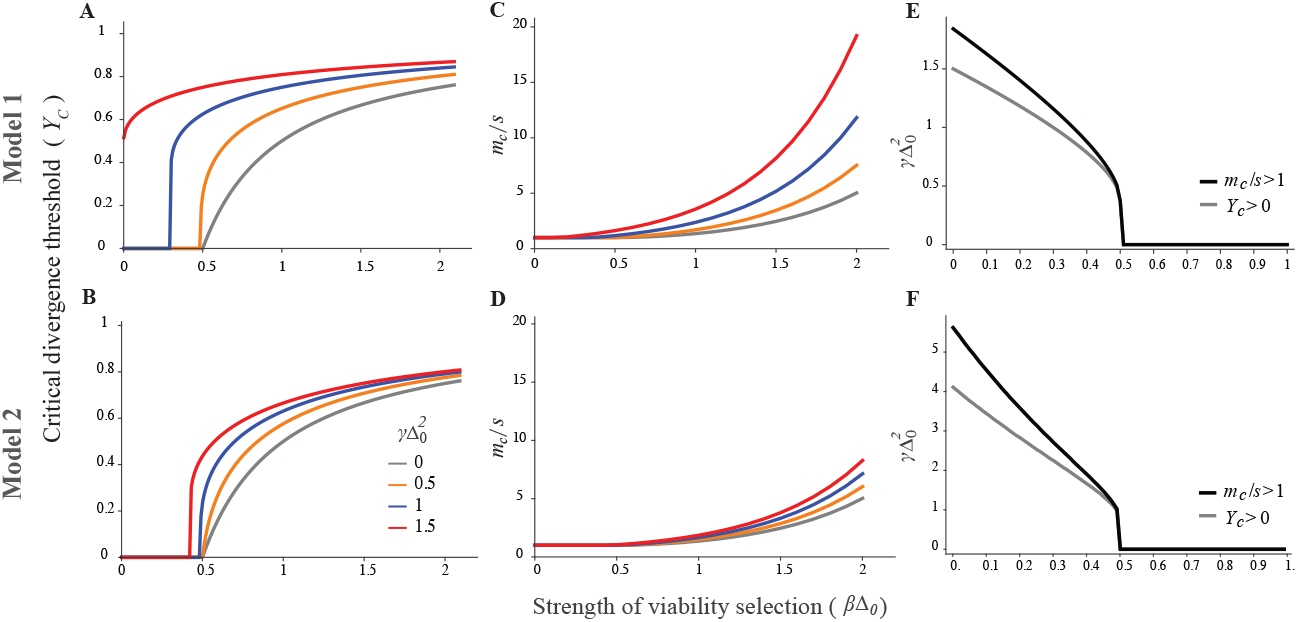
Deterministic predictions for swamping thresholds and tipping points. (A)-(B) Critical allele frequency divergence *Y*_*c*_ below which a tipping point occurs (i.e, divergence decreases sharply) vs. *β*Δ_0_, strength of viability selection. (C)-(D) Critical migration rate *m*_*c*_*/s* at which divergence is completely lost vs. *β*Δ_0_. The different colors show results for different levels of AM and sexual selection (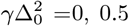,1 and 1.5). Upper panel (A, C, E) shows results for Model I (sexual selection on both sexes) and lower panel for Model II (sexual selection only on males). In panels A and B, parameter combinations with *Y*_*c*_ = 0 are those for which divergence decreases smoothly with increasing migration, i.e., there is no tipping point. In panels C and D, parameter combinations with *m*_*c*_*/s* = 1 are those for which viability and sexual selection are too weak to shift the swamping threshold above the single-locus threshold (*m/s* = 1). (E)-(F) Grey curves show the minimum level of sexual selection (as measured by 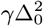) required for *Y*_*c*_ *>* 0, i.e., to generate tipping points in divergence, as a function of *β*Δ_0_, the strength of viability selection. Black curves show the minimum level of sexual selection 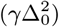 required for *m*_*c*_*/s >* 1, i.e., to shift the swamping threshold above the single-locus threshold, as a function of *β*Δ_0_. Panels E and F correspond to Model I and II respectively. All results in this figure show (large *L*) theoretical predictions obtained from eqs. 3-6.

Under sufficiently strong viability selection (*β*Δ_0_ *>* 0.5), increased swamping thresholds (*m*_*c*_*/s >* 1) emerge even in the absence of AM, as shown by the grey lines in figures 3C-3D (see also Sachdeva (2022)). Thus, in this regime, AM and sexual selection further increase the swamping threshold *m*_*c*_*/s* (beyond 1) as well as the critical level of divergence *Y*_*c*_ that can be stably maintained. This can be explained, as before, in terms of the effects of sexual selection on *m*_*e*_ and the overall genomewide barrier to gene flow. As expected, AM has a stronger effect in Model I (where sexual selection is stronger) than in Model II. For example, for *β*Δ_0_ = 1, AM with strength 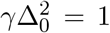 increases the critical migration threshold (*m*_*c*_*/s*) from 1.36 (with random mating) to 2.37 under Model I and 1.66 under Model II. Note also that AM makes it harder to maintain intermediate levels of divergence, with divergence becoming unstable and collapsing once mean allele frequency difference per locus approaches 0.75 under Model I and 0.66 under Model II. By contrast, under random mating (and with *β*Δ_0_ = 1), we have *Y*_*c*_ = 0.5, so that somewhat lower levels of divergence are maintained between the mainland and island, albeit at lower migration rates.

The situation is somewhat different if *β*Δ_0_ *<* 0.5; now, viability selection by itself (i.e., in the absence of AM) is too weak to generate tipping points or shift the swamping threshold *m*_*c*_*/s*. However, sufficiently strong sexual selection due to AM will lead to one or both. More specifically, given viability selection *β*Δ_0_ *<* 0.5, we can determine an assortment threshold, i.e., a minimum value of 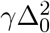 beyond which tipping points occur (*Y*_*c*_ *>* 0 in figures 3B and 3D), and a second threshold beyond which critical migration rates become higher than the single locus threshold (*m*_*c*_*/s* exceeds 1 in figures 3A and 3C); see SI, **G** for details. These assortment thresholds are depicted by grey and black curves respectively in figures 3E and 3F.

As expected, much higher levels of AM are required to generate tipping points and increase swamping thresholds when viability selection is weak, and when sexual selection acts only on males (instead of both sexes). For instance, with *β*Δ_0_ = 0.2, we have *m*_*c*_*/s >* 1 only if 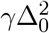 exceeds 1.41 under Model I and 3.58 under Model II (see figures 3E, 3F). However, the sex-averaged mating success of migrants relative to residents is 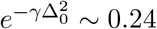 (for 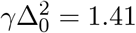) under Model I and and 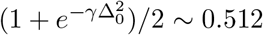 (for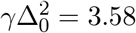) under Model II, both calculated by assuming maximum divergence between mainland and island. Thus, critical assortment thresholds are associated with much stronger sexual selection on migrants under Model I (sexual selection on both sexes) than Model II, despite AM being weaker in the former case.

### 3.3 Effect of viability and sexual selection on neutral gene flow

In the previous section, we have shown that approximations based on *m*_*e*_ accurately predict longterm adaptive divergence between the mainland and island under AM and sexual selection. In essence, divergent selection at very many loci (whether due to viability or sexual selection) results in a genome-wide reduction in gene flow, which further strengthens divergence across the genome. As discussed above, this genome-wide reduction can be quantified via *g* = *m*_*e*_*/m*, the gene flow factor at an unlinked locus, which may be viewed as a measure of RI between hybridising populations (see Westram et al. (2022)). Moreover, *g* is also related to commonly used measures of neutral differentiation such as *F*_*ST*_, which depends on both the number of migrants, *Nm*, and their reproductive value (which is equal to *g*). More concretely, at equilibrium, neutral *F*_*ST*_ between the mainland and island is: *F*_*ST*_ ∼ 1*/*(1 + 2*Nm*_*e*_) = 1*/*(1 + 2*Nm g*) on average.

We now quantify the extent to which viability and sexual selection reduce gene flow across the genome in a scenario with high migration (2*N m* = 49). Note that in the absence of divergent selection, such high levels of gene flow would erase most neutral differences between mainland and island, resulting in *F*_*ST*_ = 0.02. However, strong enough selection can maintain adaptive divergence despite high migration, thus suppressing gene flow and increasing genome-wide *F*_*ST*_. Figure 4 illustrates this by plotting *F*_*ST*_ as a function of *β*Δ_0_ and 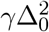 for *m/s* = 1, under the two models of AM. These figures are generated by solving for the allele frequency divergence per locus *Y* (using eq. 4) for each combination of *β*Δ_0_ and 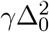,then using this to compute the gene flow factor *g* (via eq. 3), and finally computing *F*_*ST*_ = 1*/*(1 + 2*Nm g*).

**Figure 4:**
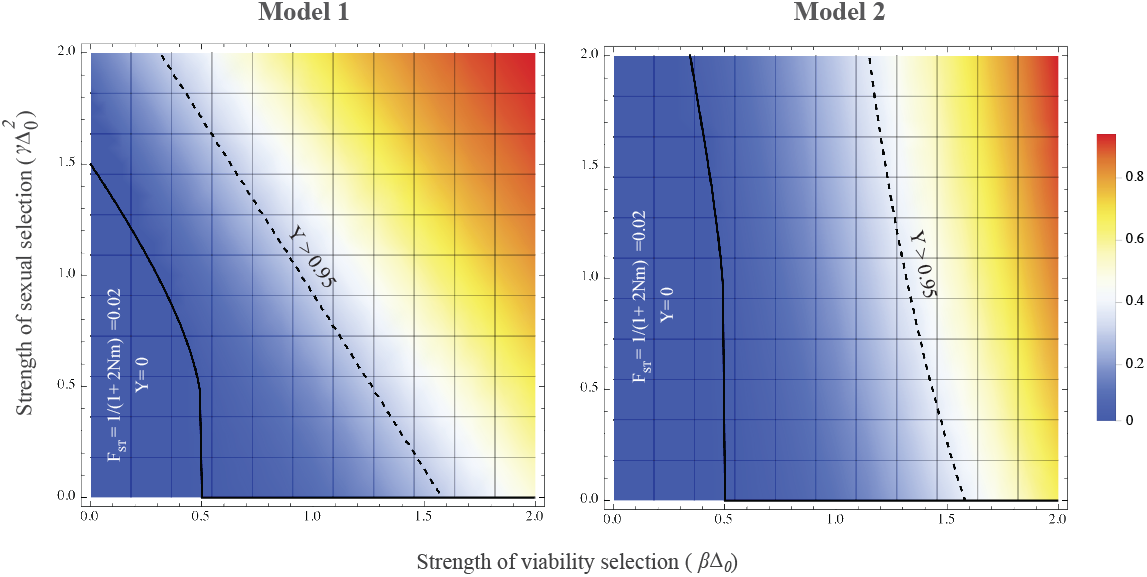
Heatmaps depicting *F*_*ST*_ at an unlinked neutral locus for different values of *β*Δ_0_ and 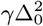 (which parameterize viability and sexual selection respectively) for Model I (sexual selection on both sexes) and Model II (sexual selection only on males) of assortative mating, assuming *m/s* = 1 and 2*Nm* = 49. The color bar on the right shows *F*_*ST*_ values ranging from *F*_*ST*_ = 0 to 1. Parameter combinations to the left of the solid line are those for which there is no allele frequency divergence (i.e., *Y* = 0) so that *F*_*ST*_ = 1*/*(1 + 2*Nm*) = 0.02; parameters to the right of this line allow for divergence, so that *F*_*ST*_ = 1*/*(1 + 2*Nm g* [Δ_0_*Y*]), where *Y* is obtained by solving eq. 4. Parameters to the right of the dashed line are those for which *Y >* 0.95: in this regime, stronger viability or sexual selection increases the genomewide barrier effect and *F*_*ST*_ without significantly affecting divergence at trait loci (which is close to its maximum possible value).

In each plot, the region to the left of the solid curve shows parameter combinations for which adaptive divergence is lost (*Y* = 0) at *m/s* = 1, so that *m*_*e*_ = *m* and 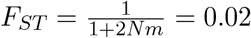. As before, increasing *β*Δ_0_ and 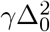 (which correspond to stronger viability and sexual selection respectively) allow non-zero adaptive divergence to be maintained, thereby reducing *m*_*e*_ and increasing *F*_*ST*_. The dashed curve in each plot depicts parameter combinations for which mean allele frequency divergence is *Y* = 0.95, which corresponds to a gene flow factor of 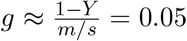 and an *F*_*ST*_ value of 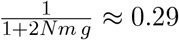 under either model of AM. To the right of this curve, i.e, with even stronger viability and/or sexual selection, there is a further increase in *F*_*ST*_, which is now *not* due to an increase in adaptive divergence (which is close to its maximum possible value). Instead higher levels of *F*_*ST*_ in this parameter regime reflect lower viability and mating success of (nearly maximally diverged) migrants, which further reduces neutral gene exchange between mainland and island.

We can also ask: do changes in the strength of viability and sexual selection multiplicatively (i.e., independently) affect gene flow (as measured by *g*), or are their combined effects stronger (or weaker) than expected from their individual effects? More concretely, consider a scenario with moderately strong viability and sexual selection 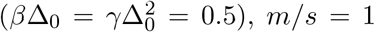,so that mean allele frequency divergence is *Y* = 0.36 (from eq. 4) and *g* = 0.64 (from eq. 3) under Model I. A 20% increase in *either* viability or sexual selection, i.e., increasing *β*Δ_0_ to 0.6 while keeping 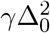 fixed at 0.5 (or vice versa) causes *g* to fall to 55% (or 74%) of its original value. Now, if viability and sexual selection were to independently reduce gene flow, then we would expect *g* to decrease by a factor of 0.55 *×* 0.74 = 0.39, i.e., to 39% of its original value in response to a 20% increase in *both* components of selection 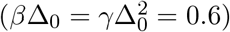.Instead, *g* decreases to 43% of its original value. Thus, in this example, the two kinds of reproductive barriers (lower survival and lower mating success of immigrants and their descendants) cause a weaker reduction in gene flow than if they were to act independently.

Such non-independent effects arise because the gene flow factor is a product of two terms– one due to viability and the other due to sexual selection (see eq. 3), both of which depend on allele frequency divergence *Y*, which in turn is influenced by both components of selection. Thus, the strongest non-multiplicative effects emerge in parameter regimes where viability and sexual selection are strong enough to significantly influence divergence levels but not so strong as to drive divergence to its maximum possible value. In the latter scenario, i.e., as *Y* approaches 1 (to the right of the dashed curves in figure 4), the effect of viability and sexual selection again becomes multiplicative.

## 4 Discussion

While it has long been recognised that sexual selection can maintain inter-specific differences, questions remain as to its importance relative to other processes during speciation (Safran et al. (2013)). Sexual selection varies significantly across species and even across environments within a species or over time (Schumer et al. (2017); Magurran and Ramnarine (2004); Rosenthal (2013)), making it difficult to identify broad patterns. Moreover, sexual selection typically acts together with other kinds of selection against heterospecifics and hybrids (e.g., due to ecological mismatch or expression of genetic incompatibilities), making it difficult to disentangle the contributions of different processes to reproductive isolation. This only underscores the broader challenges in quantifying RI– an issue that has attracted considerable attention recently (Westram et al. (2022)).

Here, we examine some of these issues in the context of a polygenic ‘magic’ trait that is under divergent ecological selection across populations, while also mediating AM and sexual selection. In such a scenario, long-term divergence depends crucially on LD between trait loci. Such LD is, in turn, a consequence of both (natural and sexual) selection and assortment, making it challenging to model. Previous analyses have relied on one of two approaches- either focusing on how trait variances are affected by selection and assortment while largely ignoring genetic details (e.g., Doebeli (1996); Kondrashov and Shpak (1998)), or explicitly modeling LD between sets of loci (Kirkpatrick (2000); Barton and De Cara (2009)), but assuming LD to be weak (to ensure tractability). Here, we employ a novel approach based on effective migration rates which bridges phenotypic descriptions (involving trait means and variances) with genic descriptions (based on divergence at individual selected sites).

An advantage of our approach is that it provides an economical description of tipping points in terms of just allele frequency differences between populations rather than a plethora of LD measures. The key idea behind our approach is that LD between introgressing alleles breaks down rapidly if these are unlinked or loosely linked (a standard assumption in quantitative genetics), allowing us to represent its effects via an effective migration rate *m*_*e*_ (or alternatively, the gene flow factor *g* = *m*_*e*_*/m*) over the longer timescales over which allele frequencies change. The gene flow factor, being the average reproductive value of migrants, is a composite measure of various processes-prezygotic and postzygotic- that affect the fitness of migrants and their descendants. In particular, it captures both the prezygotic effects of AM– which causes fewer *F*_1_ hybrids to be produced (due to sexual selection against migrants) as well as its postzygotic effects– as reflected in the lower mating success of *F*_1_s and later generation hybrids (again, due to sexual selection).

As we show here, theory based on *m*_*e*_ accurately predicts divergence across a range of parameters, including when LD is strong enough to generate so-called tipping points (Nosil et al. (2017)), but also when genetic drift is strong enough to wash away these tipping points (see SI, **D**). Moreover, it captures the consequences of sexual selection on one vs. both sexes (Model I vs. II). These models exhibit qualitatively similar behaviours– tipping points and/or shifted swamping thresholds, which, however, require much higher levels of assortment under Model II (compared to Model I) due to net (sex-averaged) sexual selection being weaker in this case. This again highlights the fact that it is sexual selection (rather than AM by itself) that influences barriers to gene flow in our model.

We also show that to good approximation, *g* is a product of two terms-one reflecting viability selection and the other sexual selection on migrants and their descendants (see eq. 3). This is in line with previous work that quantifies sexual and natural selection via probabilities of conspecific vs. heterospecific mating, or fitness of offspring from different types of mating, and then assumes that the two components of selection multiplicatively determine the total barrier to gene flow (Ramsey et al. (2003); Sobel and Chen (2014)). Note however, that in our model, the two components (due to viability and sexual selection) both depend on the existing level of divergence Δ, and thus are not independent. For instance, stronger AM and sexual selection will typically increase divergence, which will also reduce the (relative) viability of migrants and their descendants, thus further strengthening the barrier effect of viability selection, even when there is no change in the strength of viability selection. This suggests that measurements of RI are highly context and state dependent, making it difficult to extrapolate from lab measurements or between replicate hybrid zones.

Nevertheless, in theory, eq. 3 provides an unambiguous way of quantifying the contributions of natural and sexual selection to RI. In practice, this requires estimates of fitness components of individuals with different levels of migrant ancestry, which are only available in a handful of well-studied populations with multi-generational pedigrees (Bérénos et al. (2014); Chen et al. (2019)). However, such populations are necessarily spatially limited and show very little divergence or RI. More typically, we might have indirect fitness estimates for hybrids (and less commonly, for backcrosses and recombinants) from hybrid zones between divergent ecotypes. It is an open question whether combining these with experimental approaches, e.g., reciprocal transplant or mate choice experiments (that individually measure viability or mating success) might provide reliable estimates of fitness components in natural populations, thus allowing us to disentangle the contributions of natural and sexual selection to RI.

Our work also bears upon the interplay between assortative mating and sexual selection during divergence with gene flow. The distinction between the two has been highlighted in earlier work (Kirkpatrick and Nuismer (2004); Polechová and Barton (2005); Servedio and Bürger (2014)), which points out that sexual selection can reduce the mating success of outlier individuals, thus constraining genetic variance and reducing the potential for divergence (Kirkpatrick and Nuismer (2004)). By contrast, pure assortment without sexual selection always increases variance, thus facilitating divergence.

The situation is somewhat different following secondary contact between diverged populations: now, sexual selection against migrants reduces introgression of deleterious alleles. This effect is only strengthened by assortative mating (based on ancestry) which increases LD between introgressing alleles, causing them to be eliminated more effectively. In recent work, Muralidhar et al. (2022) provide a rough estimate of the effect of AM following a pulse of admixture, where an individual’s viability declines in proportion to the amount of introgressed material it carries. Their argument is as follows: the change in introgressed allele frequency per generation (due to viability and/or sexual selection) is proportional to the variance of introgressed ancestry across individuals. In an assortatively mating population with correlation coefficient *ρ* between mates (where *ρ* is a measure of the strength of AM), the variance in introgressed ancestry declines by a factor ∼ (1 + *ρ*)*/*2 per generation, *slower* than it would under random mating, allowing for more efficient selective purging of introgressing alleles.

It then follows that AM *by itself* (as a process that is distinct from sexual selection) increases net purging by a factor of 1*/*(1 − *ρ*) (Muralidhar et al. (2022)). This argument implicitly assumes that sexual selection is weak and does not cancel out the variance-increasing effect of AM; this assumption would, for instance, break down under our Model I (see also Kirkpatrick and Nuismer (2004))

The conclusions of Muralidhar et al. (2022) differ from ours– in particular, we show that divergence is shaped essentially by sexual selection on various subgroups (i.e., *F*_1_s, *BC*_1_s etc.), with little to no effect of AM (e.g., on within-subgroup variances). What might explain these differing conclusions? Muralidhar et al. (2022) analyse the consequences of a single admixture event that replaces a sizable fraction of the native population by migrants from a diverged population. In this case, individuals with recent migrant ancestry are sufficiently common to mate assortatively with one another– this causes LD between introgressing alleles to persist longer, thus allowing selection to eliminate them efficiently. By contrast, we consider ongoing migration at rate *m*, and focus on the long-term equilibrium between selection and migration. In this case, long-term adaptive divergence requires *m*_*e*_ to be lower than the typical per locus selective effect *s*, so that *m* is at most a few times larger than *s*. Thus, migration is sufficiently low that individuals with recent migrant ancestry mate predominantly with residents rather than with each other. Then, introgression depends essentially on the mating success of (or alternatively, sexual selection on) various hybrids (*F*_1_, *BC*_1_ etc.) relative to residents.

We emphasise that assuming low migration rates (*m* ≪ 1) does not imply that neutral divergence between populations is high, since the latter depends on the number of migrants *Nm* per generation, which may be large. Nor does it imply high levels of adaptive divergence (see, e.g., figure 2); indeed, this can be arbitrarily low, depending on *m/s*, and the total strength of viability and sexual selection.

The preceding discussion highlights how the role of AM (as distinct from sexual selection) depends on the context of divergence. Broadly speaking, assortment is likely to be important if diverged subgroups are present together at comparable frequencies, as in the centre of a hybrid zone. Our theoretical approximations would not apply in this case, since *F*_1_s would be sufficiently common as to mate amongst themselves and generate *F*_2_ recombinants. Moreover, many hybrid zones are characterised by a bimodal distribution of parental ancestries or equivalently, of hybrid indices (Jiggins and Mallet (2000); Bridle and Butlin (2002); Schumer et al. (2017)). The effects of AM in such a scenario are not amenable to simple descriptions which assume trait variance to increase by a constant factor (1 + *ρ*)*/*2 (Muralidhar et al. (2022)), since the correlation *ρ* between mating pairs in an assortatively mating population also depends on the distribution of phenotypic values and can be very different for unimodal vs. bimodal populations. The consequences of non-random mating in hybrid zones thus remain an important direction for future work (see also Irwin (2020)), which will require us to expand existing theoretical descriptions.

What are some limitations of our study? We have considered a model of AM via self-referencing, wherein individuals prefer mates with similar phenotypes. However, sexual selection may be less effective in suppressing gene flow if female preference and the male traits on which it acts are encoded by different genes, requiring these to be in LD. More generally, little is known about the prevalence of different mechanisms of AM and the extent to which these generate sexual selection in different ecological contexts (Safran et al. (2013); Kopp et al. (2018)), making it important to examine the robustness of theoretical findings to alternative assumptions about assortment mechanisms.

We also consider an extreme form of divergent ecological selection, where alternative alleles are favoured in the two populations at every locus. In this case, sexual selection is predominantly directional, reducing the mating success of individuals in proportion to their migrant ancestry. Alternatively, the magic trait could be under stabilizing selection towards different optima in the two populations. While selection on migrants would still be directional if trait optima are far apart, adaptive divergence can now be maintained at higher migration rates (Sachdeva and Barton (2017)). This results in a more complicated interplay between assortment and sexual selection, since mating between individuals with recent migrant ancestry becomes more common. Importantly, in this case, we need to go beyond theory that treats introgressing alleles as cascading down successively fitter genetic backgrounds, and explicitly account for F2s etc.

Finally, we neglect linkage between trait loci, which generally strengthens barriers to gene flow, at least under ecological divergent selection (Barton and Bengtsson (1986)). However, if traits are under frequency-dependent sexual selection, then linkage between trait loci may have more subtle effects (Kirkpatrick and Nuismer, 2004), even weakening barriers to gene flow under some conditions (Aubier et al., 2024). Approximations based on effective migration rates should extend to loose linkage between trait loci (Zwaenepoel et al., 2024). However, the effects of tight linkage (e.g., between alleles clustered within supergenes or inversions) are more obscure.

We focus here on the interplay between natural and sexual selection at a *fixed* level of assortment and following secondary contact between diverged populations. Other questions remain as to how assortment evolves in the first place– e.g., when mating with conspecifics might be favoured by selection, thus reinforcing divergence (Servedio and Noor (2003)). While our study does not address these questions, it does suggest that analysing the evolution of assortment in terms of the reproductive value (or effective rate of exchange) of modifiers of mate choice could be a fruitful direction for future work. More generally, effective migration rates link a ‘phenotypic’ view of RI (based on fitness components of hybrids) with a ‘genic’ view (that conceptualises RI as a process that reduces gene flow and increases divergence across the genome), making them a powerful way of understanding RI.

## Author contributions

P.S. and H.S. designed the study, did the mathematical analysis and simulations, and wrote the manuscript. The authors declare no conflict of interest.

## Acknowledgements

We thank Nick Barton for useful comments on the manuscript. This research was supported by the Scientific Service Units (SSU) of IST Austria through resources provided by Scientific Computing (SciComp).

## Data availability

Mathematica notebooks for numerical analysis and simulations are available at https://doi.org/10.15479/AT:ISTA:17344.

## Conflict of Interest

The authors declare no conflict of interest.

## SUPPLEMENTARY INFORMATION for

### A. Derivation of gene flow factor

Let *h*_*i*_ represent the fraction of individuals in the population with a migrant ancestor precisely *i* generations ago. Thus, *h*_1_ represents the fraction of *F* 1 individuals and *h*_*i*_ the fraction of (*i*−1)^*th*^ generation backcrosses (denoted as *BC*_*i*−1_). As specified in the main text, an individual with *no* migrant ancestor in the last *n* generations is designated as a ‘resident’, where *n* is an arbitrary threshold.

Thus, by definition, the fraction of residents in the population 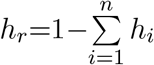. We further assume that trait values are normally distributed *within* any population subgroup: thus the distribution *P*_*r*_(*z*) of trait values among residents is normal with mean 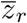 and variance *V*_*r*_; similarly, the distribution *P*_*i*_(*z*) amongst *i*^*th*^ generation descendants of migrants is normal with mean *z*_*i*_ and variance *V*_*i*_ (for any *i*). The trait value distribution across the entire population is thus the sum of *n*+1 normal distributions and is given by 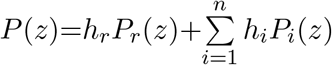. For simplicity, we will assume that the distribution of trait values amongst migrants is also normal with mean 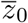 and variance *V*_0_; however, this assumption is not crucial. Finally, we will assume that *m*≪1 and perform various computations to first order in *m*. This is justified on the grounds that non-zero adaptive divergence between mainland and island requires *m* to be comparable to the typical *per locus* selection coefficient *s*, where *s*≪1 for a polygenic trait influenced by many small-effect loci.

We now consider how the proportions *h*_*i*_ and the distributions *P*_*i*_(*z*) associated with different population subgroups change within a single generation due to migration, selection and assortative mating. *Migration* from the mainland results in a fraction *m* of each subgroup being replaced by migrants.

Thus, following migration: 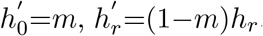,and 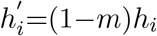 for *i*=1, … *n*. Note that migration does not change the trait distributions of existing sub-groups, so that: 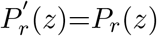 and 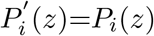.

Migration is followed by *viability selection* on the trait, wherein an individual wherein an individual with trait value *z* contributes to the pool of reproducing adults in proportion to 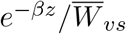.Here, 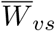 is the viability component of mean fitness and is given by: 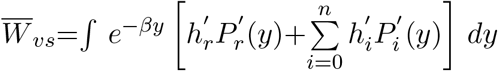. Thus, following viability selection, the proportions and trait value distributions of the different subgroups are:

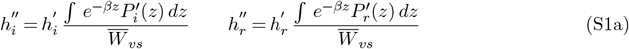

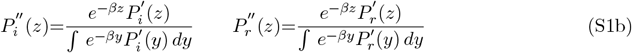

Equation S1b specifies the distribution of trait values among different population subgroups following viability selection. These distributions are still normal, but with a shifted mean 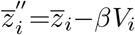 for subgroup *i* (and 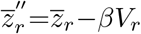 for residents). The variances within sub-groups remain unchanged (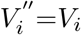 and 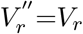)− this is because directional selection (fitness proportional to *e*^−*βz*^) acting on a (sub-)population with *normally* distributed trait values only shifts the mean without affecting the trait variance. Thus, the trait value distribution across the full population after viability selection is the sum of *n*+2 normal distributions: 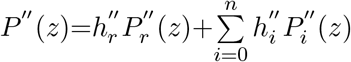. Note that this sum now includes the group ‘0’, i.e., individuals who have immigrated at the start of the generation.

In the following, we will show that *βV* is 𝒪(*m*); thus, the various *βV*_*i*_ must also be 𝒪(*m*) (assuming *V*_*i*_ to be comparable to *V*). Further, as we argue below, the various *h*_*i*_, i.e., the proportion of *F* 1s, *BC*_1_ and so on, are also 𝒪 (*m*). Thus, to first order in *m*, we can ignore the shifts in trait means (due to viability selection) within subgroups, i.e., within migrants, *F* 1s and *BC*_*i*_, since the contribution of these shifts to the shift in the overall trait mean of the full population is 𝒪 (*m*^2^). For the same reasons, we cannot ignore the shift in trait mean among residents, since 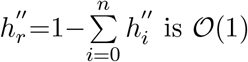.

Finally, consider the effects of *assortative mating* and *sexual selection* on population sub-groups. As described in the main text, the mating function *M* (*x, y*) specifies the probability of mating between a male and a female with trait values *x* and *y* respectively. Since an individual with a given trait value is equally likely to be male or female, the probability of mating between two randomly chosen individuals with trait values *x* and *y* is [*M* (*x, y*)+*M* (*y, x*)]*/*2. Under the weak migration (*m*≪1) assumption, migrants mate predominantly with residents to produce *F* 1s, which mate with residents to produce *BC*_1_ and so on. Further, under our definition of resident, mating between *BC*_*n*_ and residents also produces residents. Thus, after mating, we have:

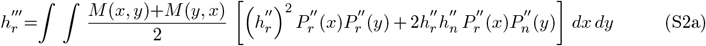

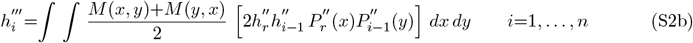

Finally, the trait value distribution among the various population sub-groups are given by:

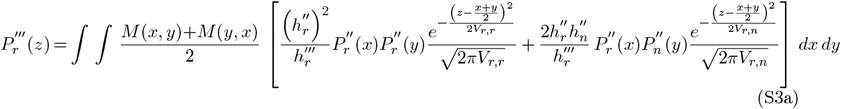

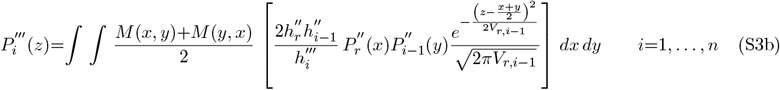

Equation S3 assumes that the trait is sufficiently polygenic that the offspring of any two individuals are normally distributed about the midparent value with a within-family variance that depends on the relatedness between parents, as in the standard infinitesimal model (Barton et al. (2017)). As discussed in the main text, we further assume that the within-family variance only depends on the subgroups to which the two parents belong. Thus, for example, the within-family variance of the offspring of any resident and *BC*_*i*_ pair is assumed to be *V*_*r,i*+1_.

At *equilibrium*, the proportions of various subgroups and their trait value distributions at the end of each generation must be the same as at the beginning, so that 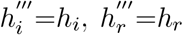,and 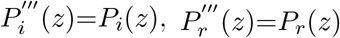,. These conditions, together with eqs. S1-S3, give us a set of equations for *P*_*i*_(*z*) and *h*_*i*_(*z*). These equations are somewhat complex, but simplify considerably if we assume that *m*≪1, and only calculate the various *h*_*i*_ to lowest order in *m*. In particular, it follows that *h*_*i*_ are 𝒪 (*m*) for all *i*. One can see this from eq. (S2b), where 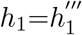 is proportional to 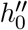 which is 𝒪 (*m*) (being the proportion of migrants in the population post viability selection), so that *h*_1_ is also 𝒪 (*m*). Similarly,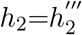 (being proportional to 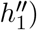 is also𝒪 (*m*), and so on. The equilibrium conditions 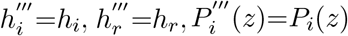 and 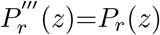 also lead to eq. 2a of the main text, which recursively relates *P*_*i*_(*z*) at equilibrium to *P*_*i*−1_(*z*). We can then use this to derive expressions for 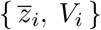, the equilibrium means and variances of trait values in each sub-group, to lowest order in *m*. In the following, we present these derivations separately for Models I and II.

Under **Model I** of assortative mating, we have:

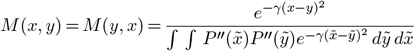

where *P* ″ (*x*) is the distribution of trait values in the population just after viability selection, which is simply: 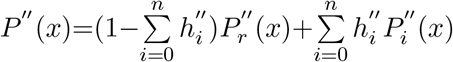. Substituting into the mating function *M* (*x, y*) and using eq. S2a, it follows that to lowest order in *m*, we have: 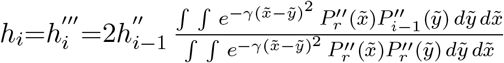 and 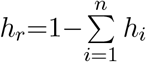 Using eq. (S1) for 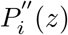 and 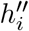 finally gives the following expressions for the equilibrium proportions of the various subgroups:

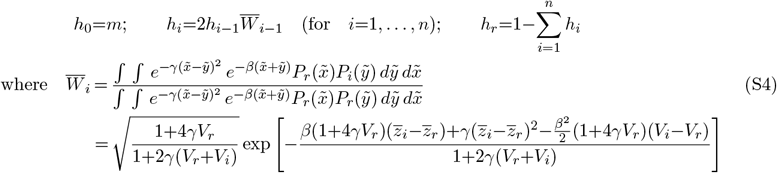

It follows from eq. (S4) above that 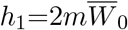 (since *h*_0_=*m*), 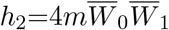, and (more generally) that: 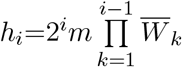. This establishes that the various *h*_*i*_ are all 𝒪 (*m*).

This establishes that the various *h*_*i*_ are all (*m*).

Equation (S4) simply states that the equilibrium proportion of *i*^*th*^ generation descendants of migrants (e.g., *BC*_1_ for *i*=2) is equal to two times the equilibrium proportions of the two parental subgroups (e.g., residents and *F* 1s for *BC*_1_s) multiplied by the mean relative fitness of the two parental subgroups.

The proportion of residents is *h*_*r*_=1−𝒪 (*m*) and their relative fitness is 1+𝒪 (*m*), whereas the various *h*_*i*_ are 𝒪 (*m*). Thus, to first order in *m*, we have: 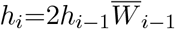,as above. Here, 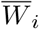 denotes the mean fitness of *i*^*th*^ generation descendants of migrants relative to residents, and is thus the same 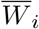 that enters the expression for the gene flow factor *g* (equation 1 of the main text). From the above equation, we see that 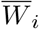 is influenced by the strength of (viability and sexual) selection as well as the trait value distributions within subgroups.

However, eq. (S4) by itself does not allow us to calculate *g*, as the means (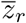 and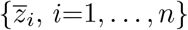) and variances (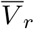 and {*V* _*i*_,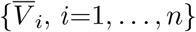) of the sub-group trait value distributions are unknown. These can be determined using eq. (S3). To do this, we will further assume that the threshold *n* in eq. (S3a) is sufficiently large that 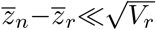,so that *P*_*r*_(*z*) (which is strictly a mixture of normal distributions; see eq. (S3a)) can be approximated by a single normal distribution with moments equal to the weighted sum of moments of the constituent distributions. Recall that *n* here is an arbitrary threshold such that an individual with *no* migrant ancestor in the last *n* generations is designated as a ‘resident’.

Also note that we are interested in deriving expressions for 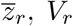 and 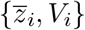 that are correct to first order in migration rate *m*. Since the effective migration rate *m*_*e*_ is simply *m* multiplied by the gene flow factor *g*, this means that we need to derive expressions for *g* or alternatively for 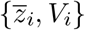) (which determine 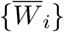,which determine *g*) that are correct to *zeroth* order in *m*. To do this, it is useful to first enumerate all the quantities that are 𝒪 (*m*). As argued above, this includes all the equilibrium proportions *h*_*i*_. Further, *βV* is also 𝒪 (*m*): this will be shown for the special case *γV* ≪1 below, but follows more generally from the fact that under migration-selection balance, the change in trait mean due to selection (which is equal to *βV*) must be balanced by the change due to migration (which is proportional to *m*). Further, since *V*_*i*_∼*V* (see below), it follows that all *βV*_*i*_ are also 𝒪 (*m*). Thus, assuming that {*h*_*i*_}, {*βV*_*i*_} and *βV* are all *O*(*m*), and using eq. (S3), we obtain the following equations for 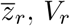,and 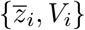 (to lowest order in *m*):

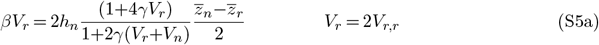

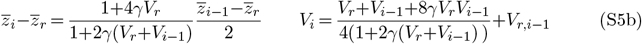

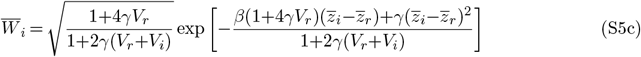

Equation (S5a) can be derived by integrating eq. (S3a) to compute the various moments of 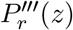,then using the equilibrium condition 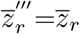 and 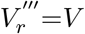,and finally retaining terms that are lowest order in *m*. Equation (S5b) follows similarly from eq. (S3b). Equation (S5c) is simply obtained from the expression for 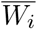 in eq. (S4) by retaining only 𝒪 (1), i.e., lowest (zeroth) order terms in *m*.

Note that eq. (S5) depends on the various segregation (within-family) variances *V*_*r,r*_ and *V*_*r,i*_, which are (so far) unknown, but must depend on allele frequencies {*p*_*i,j*_, *j*=1, …, *L*} (and {*p*_*r,j*_, *j*=1, …, *L*}) at the various trait loci in the various population sub-groups. Here, *p*_*i,j*_ (or *p*_*r,j*_) represents the allele frequency at the *j*^*th*^ locus among the *i*^*th*^ generation descendants of migrants (or among residents).

More concretely, we have: 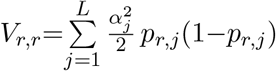 and 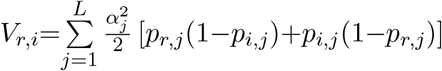. In the absence of assortment, allele frequencies among *i*^*th*^ generation descendants will simply be the average of the parental allele frequencies (i.e., of residents and (*i*−1)^*th*^ generation descendants). With assortment, {*p*_*i,j*_} are the weighted average of parental allele frequencies, which gives (under Model I): 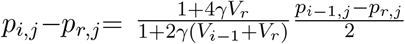 (this is analogous to equation (S5b) for 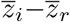). From this, we finally have:

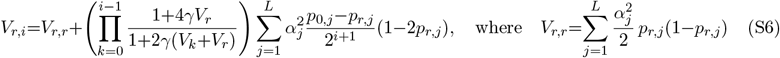

Thus, we can express all segregation variances in terms of the (as yet) unknown allele frequencies {*p*_*r,j*_} in the resident pool.

The main idea behind our theoretical approach is that if trait loci are unlinked (or loosely linked), then the allele frequency at any one locus is still (approximately) predicted by the theory for a *single* locus, but with the raw migration rate *m* replaced by an effective migration rate *m*_*e*_, which depends on allele frequencies at all the other trait loci. More concretely, in section B, we argue that at migration-selection balance (and neglecting drift), allele frequencies in the resident pool satisfy: *s*_*j*_*p*_*r,j*_(1−*p*_*r,j*_)+*m*_*e*_(*p*_0,*j*_−*p*_*r,j*_), where *s*_*j*_=*αβ*_*j*_ and *p*_0,*j*_ is the allele frequency among migrants at the *j*^*th*^ locus. If the mainland is fixed for alleles that are locally deleterious on the island (which is the scenario we consider in the main paper), we have *p*_0,*j*_=1, which yields *p*_*r,j*_=*m*_*e*_*/s*_*j*_.

We now have all the ingredients necessary for self-consistently and iteratively determining allele frequencies and mean trait divergence. The iterative procedure can be implemented as follows: one starts with an arbitrary ‘guess’ for the allele frequencies {*p*_*r,j*_}; this allows one to determine the segregation variance *V*_*r,r*_ (of offspring produced by mating between residents) using eq. (S6), as well as the trait mean 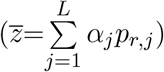 and trait variance (*V*_*r*_=2*V*_*r,r*_) among residents. These can then be used to determine the segregation variance *V*_*r*,1_ (of *F* 1 offspring produced by resident *×* migrant mating) using eq. (S6), as well as 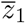 and *V*_1_ using eq. (S5b). These can now be used to determine *V*_*r*,2_ (using eq. (S6)), which can be used to determine 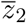 and *V*_2_ (using eq. (S5b)), allowing us to sequentially determine 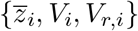 for *i*=1 … *n*. The trait means and variances can now be substituted into eq. (S5c) to obtain 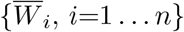,the product of which yields the gene flow factor *g*. This can be used to calculate the new *m*_*e*_, which yields the new allele frequencies via *p*_*r,j*_=*m*_*e*_*/s*_*j*_. This procedure can be iterated until the allele frequencies {*p*_*r,j*_} converge to a fixed set of values.

However, we can obtain a simple expression for *g* if *γV*_*r*_ and *γV*_*i*_ are ≪1 (see below for a discussion of this condition). Then, equation (S5) reduces to:

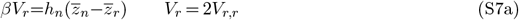

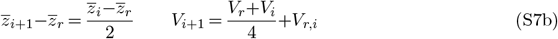

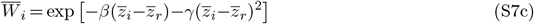

In other words, the mean phenotypic difference between each successive backcross generation and residents is half that of the previous backcross generation (eq. (S7b)), so that: 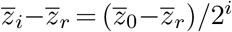. Substituting into eq. (S7c) gives 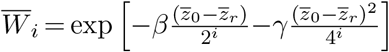, which can be substituted into 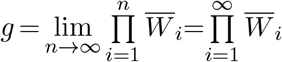. This gives the approximate expression for *g* under Model I in equation 3a of the main paper.

A second observation is that substituting 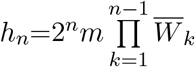 and 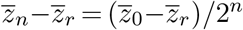 into eq. (S7a) yields 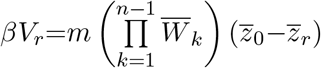. If we now take the limit *n*→∞, we have: 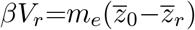.This is precisely what one would obtain by multiplying our heuristic equation *s*_*j*_*p*_*r*_(1−*p*_*r,j*_)=*m*_*e*_(*p*_0,*j*_−*p*_*r,j*_) for allele frequency dynamics by the effect size *α*_*j*_ (where *s*_*j*_=*βα*_*j*_), and summing over *j*. This demonstrates the self-consistent nature of our solution.

In summary, our simple approximation for the gene flow factor *g* (eq. 3a of the main text) follows from the more general equations (S2) and (S3) by making two additional assumptions– first, that migration rates are sufficiently low that we need only consider 𝒪 (*m*) terms, and second, that *γV*_*r*_, *γV*_*i*_≪1. As discussed above, the low migration assumption implies that: (i) the probability of mating between individuals with recent immigrant ancestry (e.g., *F* 1 *× F* 1 or *F* 1 *× BC*_1_ mating) is negligible, thus allowing us to express *g* in terms of the average fitness of successive backcrosses (see also Westram et al. (2022)), and (ii) the effect of viability selection within any sub-group of *i*^*th*^ generation descendants can be neglected, since this only has an *O*(*m*^2^) effect on the population trait mean.

We now turn to the second assumption, namely that *γV* ≪1. To understand what this implies, note that in a population with normally distributed values (with variance *V*), the phenotypic correlation between mating individuals is equal to 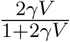 under Model I and 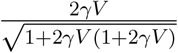 under Model II.

Thus, the correlation between mates (which can also be used as a measure of the strength of assortative mating) is ≈ 2*γV*, if *γV* ≪1, under both models. The condition that *γV*_*r*_≪1 and *γV*_*i*_≪1 can thus be interpreted as assortative mating being sufficiently weak that mate choice is ineffective and phenotypic correlation between mates is insignificant for mating *within* any subgroup. However, as long as 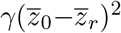 is not too small, mating *between* subgroups (e.g., between residents and *F* 1s) is still much less likely than resident*×*resident mating.

We now outline the derivation of the gene flow factor under **Model II** of assortative mating. This follows the same logic as above but involves somewhat more cumbersome expressions. Under Model II, the mating function is:

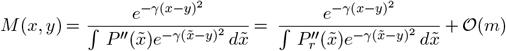

Note that in this case, *M* (*x, y*) *≠ M* (*y, x*), i.e., the probability of mating between a male and female with trait values *x* and *y* respectively is different from that of a female-male pair with values *x* and *y*.

As before, from eq. (S2), it follows that: 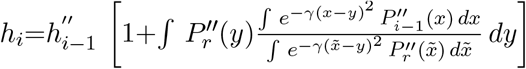 and 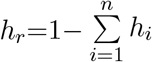 to lowest order in *m*. Using equation (S1) for 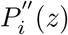 and 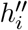 then yields the following expressions for the equilibrium proportions of the various subgroups:

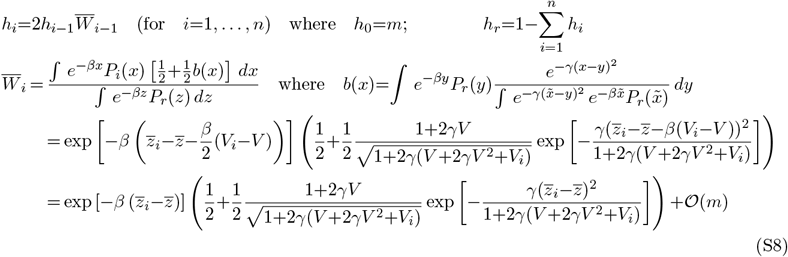

As in the case of Model I, this yields: 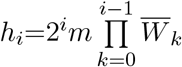

We can now use eq. (S3) together with the function *M* (*x, y*) for Model II (as given on the previous page) to derive expressions relating 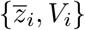 and 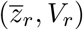 to each other, restricting ourselves (as before) to computations that are correct to lowest order in *m*. However, there is now an important difference (as compared to Model I): under Model II, if trait values are normally distributed among residents and (say) among *F* 1s, then the distribution of trait values among their (*BC*_1_) offspring will be the *sum* of two different normal distributions– one corresponding to offspring of resident father and F1 mothers and the other to offspring of resident mothers and F1 fathers. These distributions may be slightly shifted away from the midparent value towards the F1 mean (for offspring of F1 mothers and resident fathers) or towards the resident mean (for offspring of resident mothers and F1 fathers, and provided 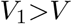) and thus are not identical. However, one can show that the squared difference between the mean of these two distributions relative to the variance is proportional to *γV*, and will thus be small for *γV* ≪1. Then, we can approximate the mixture of two normal distributions by a single normal distribution with moments equal to the weighted sum of the constituent distributions. This is the additional assumption that we need to make in order to derive expressions for 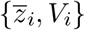 and 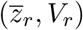 under Model II (in analogy to eq. (S5) for Model I).

However, these expressions are somewhat cumbersome and we do not provide them here. Instead we focus on a parameter regime in which *γV*_*r*_, *γV*_*i*_ are ≪1. Then, as in Model I, the expressions for the trait means and variances and average relative fitness of the subgroups simplify considerably, and we have:

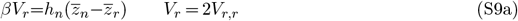

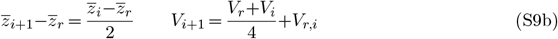

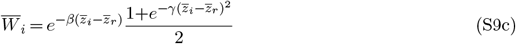

Note that equations (S9a) and (S9b) are identical to eqs. (S7a) and (S7b). However, the mean fitness of the *i*^*th*^ subgroup is different under the two models (eq. (S7c) vs. eq. (S9c)), reflecting the fact that mating success is reduced for both male and female non-resident individuals under Model I but only for non-resident males under Model II. It follows from eq. (S9b) that 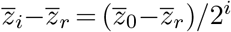.Substituting into eq. (S9c) gives: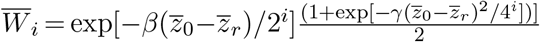, which can be substituted into 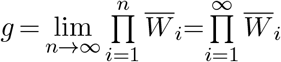.This gives the approximate expression for *g* under Model II (equation 3b of the main paper).

### B. Deterministic allele frequency dynamics at a trait locus under Linkage Equilibrium (LE)

In this section, we derive an expression for *direct* selection at a trait locus by assuming Linkage Equilibrium (LE), i.e., by neglecting statistical associations or LD between loci. The key result here is that sexual selection contributes negligibly to direct selection at any trait locus if the trait is sufficiently polygenic (see below for a more precise statement). However, as we show in section A, sexual selection does reduce the gene flow factor, *g* (or *m*_*e*_ = *m g*), which accounts for the effects of LD between deleterious alleles (which we ignore in this section).

Consider the change in allele frequency (Δ*p*)_*sel*_ due to a single generation of natural and sexual selection at a focal locus with additive effect *α* and allele frequency *p*. This can be expressed as:

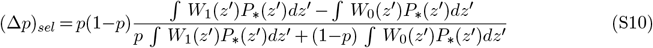

where *W*_1_(*z*′) (or *W*_0_(*z*′)) is the fitness of an individual that carries the ‘1’ (or ‘0’) allele at the focal locus and has trait value *z*=*z*′+*α* (or *z*=*z*′). Thus, *z*′ is the contribution to trait value due to all loci minus the focal locus, and *P*_∗_(*z*′) is the distribution of *z*′ in the population. Note that under the LE assumption, this distribution must be the same for individuals carrying the ‘0’ vs. ‘1’ allele at the focal locus. Now, if *L* is large, then *P*_∗_(*z*′) is approximately normal with mean 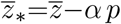 and variance *V*_∗_=*V* −*α*^2^*p*(1−*p*), where *z* and *V* are the mean and variance of trait values (as determined by all *L* loci) in the population.

Under **Model I**, an individual with trait value *z* has fitness 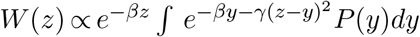, where 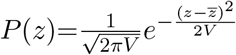 is the distribution of trait values in the population. This gives: 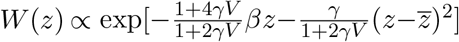Now substituting *W*_1_(*z*′)=*W* (*z*′+*α*) and *W*_0_(*z*′)=*W* (*z*′) into eq. (S10), performing the integrals over the (normal) distribution *P*_∗_(*z*′), and Taylor expanding in powers of *α*, one obtains:

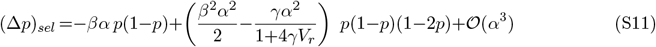

Under **Model II**, the fitness of an individual with trait value *z* is: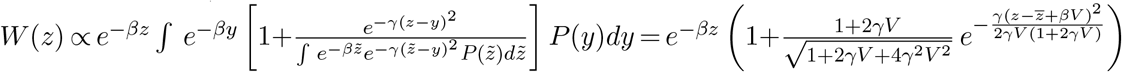. As before, we can substitute *W*_1_(*z*′)=*W* (*z*′+*α*) and *W*_0_(*z*′)=*W* (*z*′) into eq. (S10), integrate over *P*_∗_(*z*′), and Taylor expand in powers of *α*. This gives:

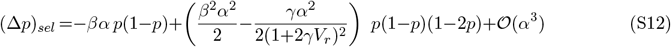

Thus, if *α* is sufficiently small (see below for a more precise condition), then to leading order in *α*, we have: (Δ*p*)_*sel*_ ≈−*βα p*(1−*p*) under both models of assortative mating. If we now assume the mainland to be fixed for alleles that are locally deleterious on the island, then the per generation change in allele frequency due to migration is simply (Δ*p*)_*mig*_=*m*(1−*p*). Then, at migration-selection balance (and assuming LE): Δ*p*=(Δ*p*)_*sel*_+(Δ*p*)_*mig*_=−*sp*(1−*p*)+*m*(1−*p*)=0, where *s*=*βα*. Thus, the equilibrium allele frequency is *p*=*m/s* if *m<s*, and *p*=1 otherwise.

How small does *α* need to be for us to be able to neglect *𝒪*(*α*^2^) terms in equations (S11) and (S12)? This requires that both *β*^2^*α*^2^ and *γα*^2^ be much smaller than *βα*, which in turn requires: *βα*≪1 and *α≪β/γ*. The first condition, namely, *s*=*βα*≪1 simply means that the change in allele frequency in any generation due to selection (or migration) should be sufficiently small that allele frequency dynamics can be approximated by continuous time equations (as is standard in population genetics).

The second condition *α≪β/γ* can be rewritten as 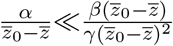 : this translates into the requirement that the relative contribution of any locus to mean trait divergence is much smaller than the ratio of the sexual selection component of migration load relative to the viability selection component. This condition is satisfied as long as the two components are comparable in magnitude.

### C. Deterministic simulations under the hypergeometric model

We perform deterministic simulations of the mainland-island model based on the hypergeometric model. The order of events on the island is migration, viability selection, assortative mating (which generates sexual selection), and free recombination between parental genotypes. The mainland is assumed to be fixed for the alleles that are locally deleterious on the island. Additionally, unless specified otherwise, the island is initially fixed for the favorable allele at all trait loci, as in a secondary contact scenario.

Let *P* (*z*) be the probability distribution of trait values 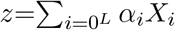 on the island, where *α*_*i*_ is the effect size and *X*_*i*_=0, 1 denote alternate alleles at locus *i*. Under the hypergeometric model, all *L* loci have equal effect (denoted by *α*) on the trait and all multi-locus genotypes corresponding to a given trait value are equally abundant in the population. Thus, we can express the trait value of an individual as *z*=*αk*, where *k* is the number of ‘1’ alleles carried by the individual. Further, the distribution *P* (*z*) is equivalent to the distribution *P* (*k*) on the island. Starting with a distribution *P*_*t*_(*k*) at the end of generation *t*, the distribution is evolved under migration, viability selection and assortative mating as follows: :

### Migration

*P*′(*k*)=(1−*m*)*P*_*t*_(*k*)+*mδ*_*k,L*_, where *δ*_*k,L*_=1 if *k*=*L* and is 0 otherwise.

### Viability selection

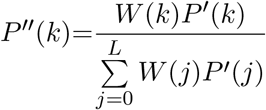,where 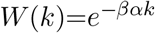

### Assortative mating and free recombination

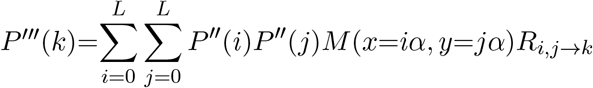. Here, *R*_*i,j*→*k*_ is the probability that parents carrying *i* and *j* alleles (say, of type ‘1’) produce an offspring with *k* ‘1’ alleles. This is given by: 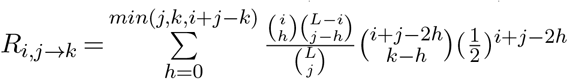, where *j*≤*i*, and *max*(0, *i*+*j*−*L*) *<k <min*(*i*+*j, L*), and *L* is the number of loci (see equation A2 in Barton (1992)). Also, *M* (*x, y*) is the probability of mating between a male and female with trait values *x* and *y* respectively, where *x*=*iα* and *y*=*jα* (see section A for concrete formulae under Models I and II). Thus, at the end of the (*t*+1)^*th*^ generation, we have: *P*_*t*+1_(*k*)=*P* ^‴^ (*k*).

The steps above are iterated in each generation until equilibrium is attained, i.e., the average allele frequency 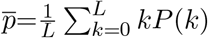 (averaged over all trait loci) no longer changes with time. In the following, we will depict our results in terms of the average allele frequency divergence per trait locus, 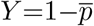.

### D. Accounting for the effects of genetic drift on divergence

In the main paper, we focus on the deterministic limit of the model. However, our analytical approach generalises to finite populations, where we can obtain the expected trait divergence by integrating over the *distribution* of allele frequencies on the island. The distribution of allele frequencies at any locus is assumed to be governed by the balance between mutation, selection, genetic drift and *effective* migration (eq. 5 of the main paper), where the last depends on the expected trait divergence between mainland and island (eq. 3, main paper), allowing for a self-consistent solution. We now test the validity of our finite population approximations by comparing against individual-based simulations.

These simulations track *N* haploid individuals on the island, each carrying *L* unlinked, biallelic trait loci. At the start of the simulation, the island is perfectly adapted, with all individuals carrying the locally adaptive allele at all *L* loci. In each generation, we choose a Poisson-distributed number of individuals (with mean *Nm*), and replace them with mainland individuals that carry the locally deleterious allele at each trait locus. Migration is followed by mutation, where the allelic state at each locus in each individual is flipped independently with probability *µ*. Mutation is followed by selection of parents, which is performed differently under the two models. In Model I, we sample (with replacement) *N pairs* of parents with sampling weights proportional to the relative fitness of the pair, which is: 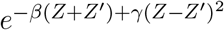 for a pair of individuals with trait values *Z* and *Z*′. In Model II, we first sample (with replacement) *N* individuals in the *female role*, i.e., with sampling weights proportional to female fitness, which is: *e*^−*βZ*^ for an individual with trait value *Z*. For each such individual (having, say, trait value *Z*′), we then choose a mate by sampling individuals (with replacement) with weights proportional to male fitness, which is: 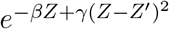 for an individual with trait value *Z*. Thus, in both Models I and II, we have *N* parental pairs at the end of the sampling step. Haploid offspring are then created by free recombination between pairs, i.e., by sampling alleles from one or other parent with probability 1*/*2 independently at each locus. These steps are iterated over successive generations, until the mean and variance of the trait value distribution on the island stop changing in time.

**Figure S1:**
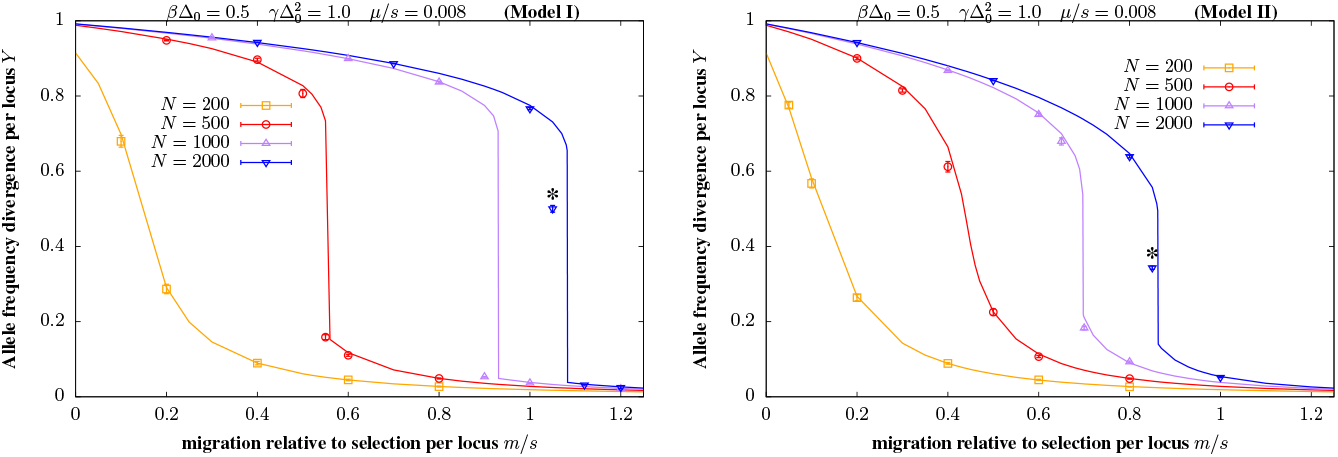
Comparison of individual-based simulations with theoretical predictions. Left and right panels show *Y*, the mean allele frequency divergence per locus as a function of *m/s*, the migration rate scaled by per locus selection coefficient, under models I and II of assortative mating. The magic trait is encoded by *L*=80 unlinked loci, with *βα*=*s*=6.25*×*10^−3^ and *γα*^2^=1.5625*×*10^−4^, so that the scaled strengths of viability and sexual selection are respectively: *β*Δ_0_=*βαL*=0.5 and 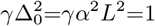.Each locus undergoes bidirectional mutation at rate *µ*=5*×*10^−5^ per generation, so that *µ/s*=0.008. The different colors depict results for different population sizes *N*. Lines show theoretical predictions (obtained by numerically solving equation 5 of the main text); symbols show results of individual-based simulations. Each simulation point is obtained by averaging over allele frequencies at every 40^*th*^ generation over a period of 5000 generations (after equilibration) in each simulation, and further averaging this over 20 simulation replicates; error bars depict standard error across replicates. The two points marked by ∗ have not equilibrated in the run time of the simulation, with the equilibrium values likely to be lower than what is depicted in the figures.

Figure S1 shows allele frequency divergence per locus as a function of *m/s*, for various values of *N*, for moderately strong viability and sexual selection (*β*Δ_0_=0.5,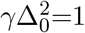) under the two models of AM. Symbols depict results of individual-based simulations, while lines show theoretical predictions obtained by self-consistently solving equation 5 of the main text, while using equation 3 of the main text to approximate the gene flow factor.

Three comments about figure S1 are in order. First, we find that theoretical approximations that account for drift are in fairly good agreement with simulation results across an order of magnitude of *N* (and *Ns*) values. Modest deviations between theory and simulations can be observed close to tipping points (as are also seen in the deterministic case; see figure 2 of the main text)– these are likely due to the relatively crude approximation that we use for the gene flow factor. Second, as expected, smaller populations are swamped at lower levels of gene flow. This is true even in the absence of multi-locus barriers, with local adaptation at a single divergently selected locus being appreciably reduced if *Ns* ≲ 20, for (say) *m/s*=0.5. Third, tipping points (characterised by an abrupt collapse in divergence) can only be observed for sufficiently large *Ns* (*N* ≥500 or *Ns*≥3.125 in Model I and *N* ≥1000 or *Ns*≥6.25 in Model II). In smaller populations (i.e., with smaller *Ns*), divergence decays smoothly with increasing gene flow under both models. The absence of tipping points in small populations is explained by the fact that the positive feedback between increasing adaptation at individual loci and declining *m*_*e*_ across the genome (which is ultimately responsible for tipping points) is weaker when evolutionary dynamics at any locus are dominated by genetic drift.

### E. Dependence of divergence per locus on the number of loci (deterministic predictions)

The theoretical predictions in equations 3 and 4 of the main paper (see also section A) are independent of *L*, the number of trait loci, and depend on only three composite parameters— the rate of migration relative to per locus selection *m/s* (where *s*=*βα*), the strength of viability selection on migrants (as quantified by *β*Δ_0_) and the strength of sexual selection on migrants (as quantified by 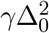). These predictions are correct to first order in small parameters *m, s* etc., and are thus expected to become increasingly inaccurate as selection per locus becomes stronger. In particular, for a given strength of viability selection *β*Δ_0_ and sexual selection 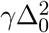 (where Δ_0_=*αL* is the maximum possible trait divergence between mainland and island), we expect our predictions to become more accurate as we consider traits determined by larger and larger number of loci of weaker effect, i.e., as we approach the limit *α*→0, *L*→∞ with Δ_0_=*αL* held constant.

We now verify these expectations by comparing theoretical predictions with the results of deterministic simulations of traits influenced by *L*=10, 40, 100 loci, for a specific strength of viability and sexual selection 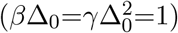 for the two models of assortative mating. In order to keep the maximum possible divergence Δ_0_=*αL* (as well as *β*Δ_0_ and 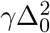) constant, we simultaneously decrease *α* while increasing *L*. For small *L*, say *L*=10, the effect of LD at any locus is weaker than what is predicted by our expressions, so that divergence is lost at lower values of *m/s*. However, as expected, there is better agreement between simulations and theory with increasing *L*, so that *m*_*c*_*/s* for a trait with *L*=100 loci only slightly lower than the analytical prediction (compare purple lines with black line in figure S2). More generally, we observe larger deviations from theory when *β*Δ_0_ and/or 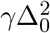, *L* are large. Thus, if net selection is strong, then *L* must be correspondingly large to observe good agreement between theory and simulations.

**Figure S2:**
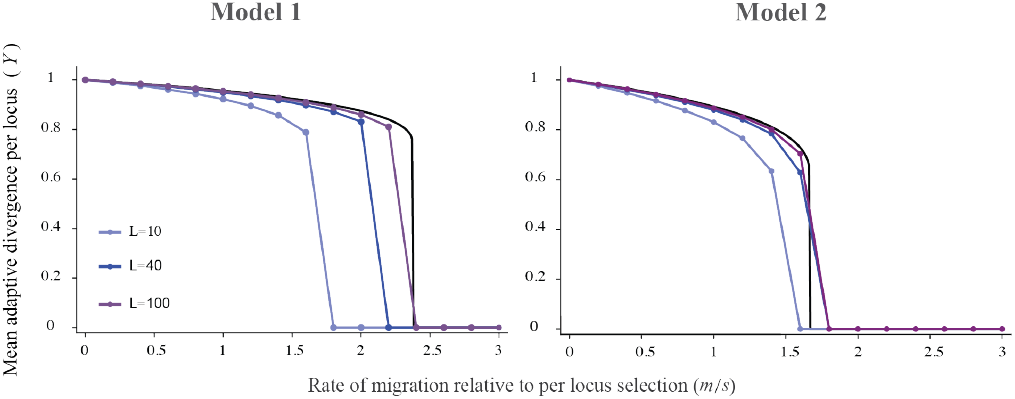
Mean allele frequency divergence per locus (Y) between mainland and island vs. migration rate relative to per locus selection (*m/s*), for Models I and II of assortative mating, with 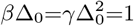 in both panels. Different colours show the results from the deterministic simulation with *L*=10, 40, and 100 loci, where *α* is decreased as *L* is increased to keep Δ_0_=*αL* constant. The black curve shows the analytical (large *L* or, equivalently, small *s*) predictions for the two models, obtained by numerically solving eq. 4 in the main text.

### F Dependence of equilibrium divergence on initial divergence levels (deterministic predictions)

Throughout the main text, we have considered a secondary contact scenario with high initial divergence between mainland and island. However, under intermediate or strong assortment, where two stable equilibria are possible (marked by crosses in figure S4B and C), the island population can evolve towards either equilibrium under gene flow, depending on initial conditions. More specifically, starting with high initial divergence (*Y* ∼1), the population approaches the stable equilibrium associated with the higher level of divergence, provided *m<m*_*c*_. By contrast, if *Y* is initially low, the population approaches the low (or zero) divergence equilibrium. In other words, in this regime, gene flow is strong enough to prevent genetically similar populations from diverging (despite divergent selection), but not strong enough to erode divergence in populations that are already strongly diverged.

**Figure S3:**
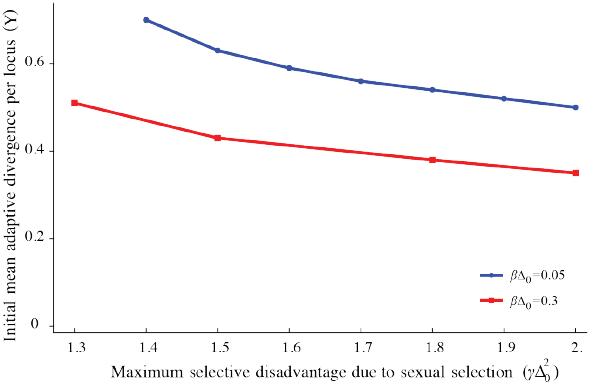
Minimum initial allele frequency divergence per locus required for populations to evolve towards the high-divergence equilibrium, plotted against the strength of sexual selection 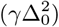 for Model I of assortative mating. Different colours show results for two different strengths of viability selection *β*Δ_0_=0.05 and 0.3. All results are obtained from deterministic simulations with *L*=40 loci.

Figure S3 explores how equilibrium divergence depends on initial divergence using deterministic simulations initialised with different values of *Y*, the mean per locus allele frequency divergence. Note that under the hypergeometric model (where all trait loci are exchangeable), this amounts to assuming an allele frequency divergence *Y* at each locus. The population is then allowed to equilibrate under gene flow. We consider assortment levels at which alternative stable equilibria exist, so that populations can evolve towards either equilibrium, depending on the initial divergence *Y*. Figure S3 shows the minimum initial divergence per locus required for populations to attain the ‘high-divergence’ equilibrium, as a function of 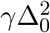, the strength of sexual selection, for two different values of 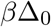. As expected, long-term divergence can be maintained more easily, i.e., even with moderate levels of initial divergence, if net (viability or sexual) selection is strong.

### G Critical divergence thresholds and migration rates (deterministic predictions)

As described in the main text (see also section B), the dynamics of the average allele frequency divergence per locus, 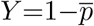 can be approximated as:

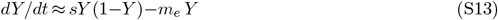

This is the same as the equation for a single selected locus, but with *m* replaced by the effective migration *m*_*e*_, which captures the effect of LD between the focal locus and all other trait loci. Furthermore, *m*_*e*_=*m g*(*Y*), where *g* is the gene flow factor (given by equation 3 in the main text), which itself depends on the average divergence level (and on *β*Δ_0_ and 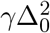,which quantify respectively the strength of viability and sexual selection). Thus, at migration-selection-assortment equilibrium, we have *dY/dt*=0, so that the equilibrium divergence level is given by the solution(s) of:

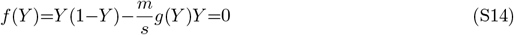

This yields one or more equilibria *Y* ^∗^, which are stable provided *f* ′(*Y* ^∗^)*<*0. This can be visualised by plotting *f* (*Y*) vs *Y* (see figure S4). For any *m/s*, the point at which the curve intersects the horizontal axis corresponds to an equilibrium; it is stable if the curve is downward-sloping at that point and unstable otherwise. *Y* ^∗^=0 is always an equilibrium and is stable if *m/s>*1. Additionally, there may be other equilibria with *Y* ^∗^*>*0 which satisfy 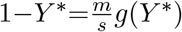.

For weak selection, say when 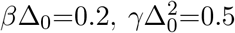 (see figure S4A), there is an unstable equilibrium at 0 and a non-zero stable equilibrium (as marked by a cross) when *m/s<*1. Increasing *m/s* causes this equilibrium to shift to smaller values; in other words, there is a smooth decline in adaptive allele frequencies until divergence is lost at the critical migration threshold, *m*_*c*_*/s*=1. For *m/s*≥1, the only stable equilibrium is at *Y* ^∗^=0.

The behaviour of equilibria is qualitatively different if the strength of (viability and/or sexual) selection is intermediate or high. For example, for 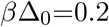 and 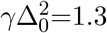 (figure S4B), there is a single non-zero stable equilibrium at low *m/s*, as with weak selection. Increasing *m/s*, say to 0.93 gives rise to two stable non-zero equilibria (marked as crosses), separated by an unstable equilibrium (in addition to an unstable equilibrium at 0). As *m/s* increases further (to 0.949), the stable ‘high-divergence’ equilibrium collides with the unstable equilibrium at a critical divergence threshold *Y*_*c*_ (marked by a circle). A further increase in *m/s* causes this high-divergence equilibrium to vanish, so that only the alternative ‘low-divergence’ stable equilibrium remains. We refer to this abrupt collapse of the high-divergence equilibrium as a ‘tipping point’. A further increase in *m/s* causes the low-divergence equilibrium to approach 0, until *Y* ^∗^=0 becomes the only stable equilibrium at *m*_*c*_*/s*≥1. Thus, with intermediate selection, we observe a tipping point (at which divergence collapses abruptly from high to low but non-zero levels) at a migration threshold *m*^∗^*/s* that is slightly below *m*_*c*_*/s*=1, which is the critical migration threshold for complete loss of divergence (blue curve in figure S5).

**Figure S4:**
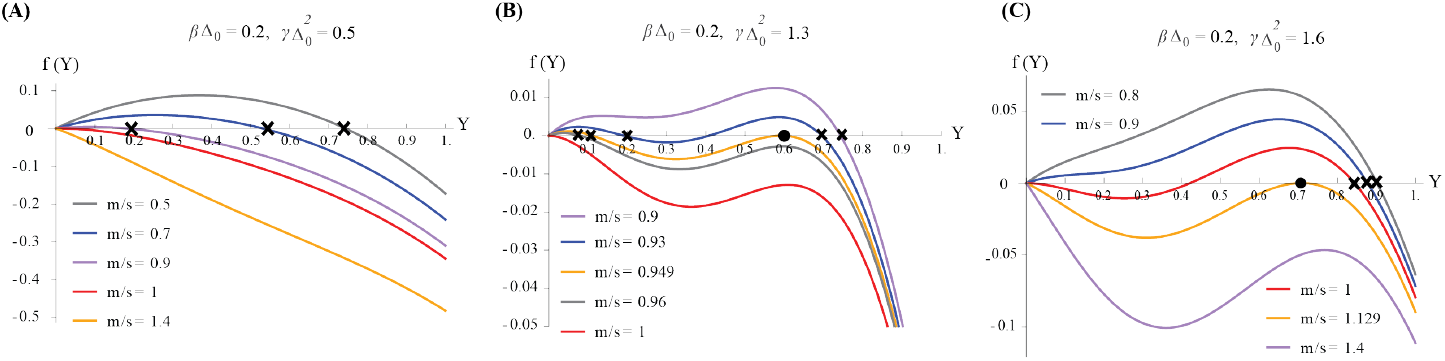
Different behaviours of the equilibria are visualized by plotting *f* (*Y*) vs. *Y* for Model I of assortative mating, where *Y* is the mean allele frequency divergence per locus and 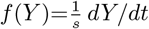 is given by equation S14. For a given *m/s*, equilibria *Y* ^∗^ are the points of the curve that intersect the horizontal axis, i.e., for which *f* (*Y* ^∗^)=0, and are stable if the curve is downward sloping, i.e., if *f* ^*i*^(*Y* ^∗^)*<*0. (A) Weak selection 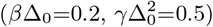:there is a non-zero stable equilibrium (marked by a cross) if *m/s<*1, which disappears at a critical migration threshold, *m*_*c*_*/s*=1, beyond which 0 is the only stable equilibrium. (B) Intermediate selection 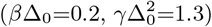: there are two non-zero stable equilibria for certain migration thresholds, e.g., for *m/s*=0.93 (see crosses), separated by an unstable equilibrium. With increasing *m/s*, the unstable and stable equilibria come closer and collide at a critical divergence threshold *Y*_*c*_ (marked by a circle for *m/s*=0.949). This equilibrium vanishes with a slight increase in *m/s*, say when *m/s*=0.95, resulting in a sudden drop in divergence to a low but non-zero value (marked by a cross). A further increase in *m* causes divergence to drop gradually until 0 becomes the only stable equilibrium beyond the critical migration threshold *m*_*c*_*/s*=1. (C) Strong selection 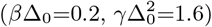:As with weak and intermediate selection, there is one non-zero stable equilibrium when *m/s<*1 and 0 is an unstable equilibrium. When *m/s*≥1, there are two stable equilibria– one at 0, and the other at a high divergence level, separated by an unstable equilibrium. At a critical migration threshold *m*_*c*_*/s*=1.129, the stable and unstable equilibria collide– this occurs at a divergence level *Y*_*c*_ (marked by a circle). Beyond this critical migration threshold, 0 is the only stable equilibrium. Note that the curve *f* (*Y*) always has a maximum at *Y* =*Y*_*c*_ (marked by circles in 1B and 1C).

With higher selection, say for *β*Δ_0_=0.2 and 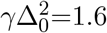 (see figure S4C), there is a single stable high-divergence equilibrium for *m/s<*1. As *m/s* crosses 1, the equilibrium *Y* ^∗^=0 also becomes stable, so that there are now two stable equilibria– the zero divergence and the high-divergence equilibrium separated by an unstable equilibrium. A further increase in *m/s* causes the unstable and stable (high-divergence) equilibria to approach each other, until at a critical migration threshold (here, *m*_*c*_*/s*=1.129), the two equilibria collide– this occurs at a critical divergence value *Y*_*c*_ (marked by a circle). Beyond this migration threshold, *Y* ^∗^=0 becomes the only stable equilibrium. Thus, in the strong selection regime, there is a critical migration threshold (*m*_*c*_*/s>*1) at which a tipping point occurs such that divergence collapses abruptly from a high level to zero (figure S5).

Below, we use equation S14 to derive the divergence threshold (*Y*_*c*_) beyond which there occurs a tipping point or sudden drop in divergence, and the critical migration threshold *m*_*c*_*/s*, beyond which divergence goes to zero. From figures S4B and S4C, we can see that the critical divergence threshold *Y*_*c*_ (at which the high-divergence equilibrium suddenly vanishes) has the property that *f* (*Y*_*c*_)=0 and *f* ^′^(*Y*_*c*_)=0. This gives:

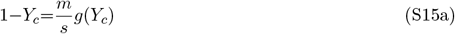

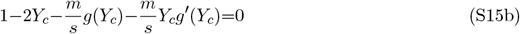

**Figure S5:**
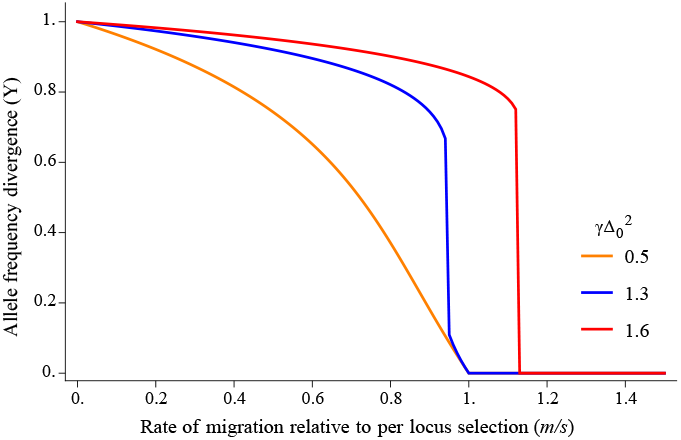
Allele frequency divergence *Y* vs. rate of migration relative to per locus selection (*m/s*) for a given strength of viability selection (*β*Δ_0_=0.2) and varying strengths of sexual selection (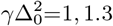and 1.6). The plot illustrates different possible behaviours associated with loss of local adaptation: with moderate sexual selection (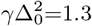;blue curve), the tipping point occurs at a migration rate lower than *m*_*c*_*/s*=1 (at which divergence is completely lost). With stronger sexual 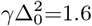;red curve), the tipping point and loss of divergence occur at the same threshold *m*_*c*_*/s>*1.

Note that *m/s* in equation S15 is the migration threshold associated with *Y*_*c*_. This migration threshold is less than *m*_*c*_*/s* (which is equal to 1) when assortment is weak or moderate (see figure S4B) and the same as *m*_*c*_*/s* under strong assortment (see figure S4C).

#### Random mating

With no assortment (*γ*=0), we have: 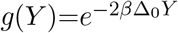,so that *g* ′ (*Y*_*c*_)=−2*β*Δ_0_ *g*(*Y*_*c*_), which when substituted into eq. S15 gives: 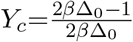.

#### Model I of assortative mating

In this case, we have: 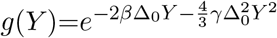,so that 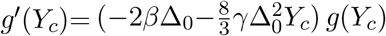. Substituting this into eq. S15b, and using eq. S15a gives:

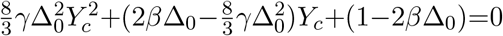. This can be solved to get two roots for *Y*_*c*_ of which

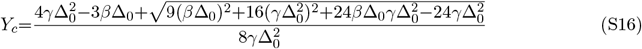

is the biologically meaningful solution.

#### Model II of assortative mating

In this case, we have: 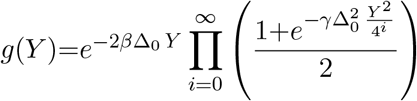,so that 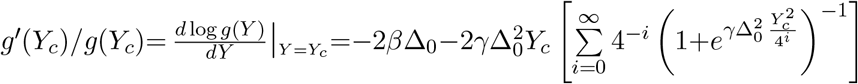. Substituting this into eq. S15b and using eq. S15a gives the following equation for *Y*_*c*_:

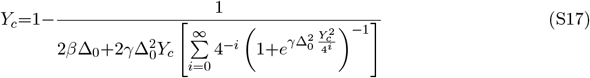

which is solved numerically by approximating the infinite sum by the first 20 terms (which is sufficient for convergence).

For both models of assortative mating, *Y*_*c*_ can be substituted in equation S15a to get *m*^∗^*/s*, which is the migration threshold associated with the tipping point. This is the same as *m*_*c*_*/s* (at which divergence is completely lost) under strong selection (e.g., in figure S4C). However, under more moderate selection, tipping points occur at *m*_∗_*/s <m*_*c*_*/s*, where *m*_*c*_*/s*=1 (e.g., in figure S4B). Thus, we have: *m*_*c*_*/s*= Max[1, *m*^∗^*/s*]. These predictions for *Y*_*c*_ and *m*_*c*_*/s* are depicted in figure 3A-D of the main text.

We can also use the above predictions for *Y*_*c*_ and *m*_*c*_*/s* to compute the minimum strength of sexual selection required for: (a) tipping points in divergence (associated with a non-zero *Y*_*c*_), and (b) shifted critical migration thresholds, i.e., *m*_*c*_*/s>*1, given a certain strength of viability selection. These predictions are shown in figure 3E-F of the main text. With random mating, *Y*_*c*_*>*0 requires *β*Δ_0_*>*1*/*2 and is also associated with *m*_*c*_*/s>*1 (also see Sachdeva (2022)). Put another way, if the strength of viability selection exceeds *β*Δ_0_=1*/*2, then we have *Y*_*c*_*>*0 and *m*_*c*_*/s>*1 even with zero sexual selection. Thus, we will focus on the parameter regime with *β*Δ_0_*<*1*/*2.

Under Model I of assortative mating, *Y*_*c*_*>*0 requires that the term under the square root in equation S16 be positive. This gives the minimum strength of sexual selection required to observe a tipping point (given *β*Δ_0_) as:

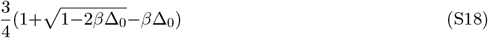

To find the minimum strength of sexual selection required to shift the critical migration threshold beyond 1, we substitute *m/s*=1 into eq. S15a, which gives *g*(*Y*_*c*_)=1−*Y*_*c*_. If we now substitute the expression for *Y*_*c*_ under Model I (as given by eq. S16), then we obtain an equation that only involves *β*Δ_0_ and 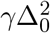.This can be solved numerically to obtain the value of 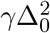 at which *m*_*c*_*/s* just exceeds 1, for any given value of *β*Δ_0_. Obtaining similar predictions under Model II is non-trivial. Thus, in this case, we numerically solve equation S17 to find the minimum value of 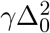 at which *Y*_*c*_*>*0 and similarly for *m*_*c*_*/s>*1. These predictions are depicted in figure 3E-F of the main text.

## Notes

https://doi.org/10.15479/AT:ISTA:17344

### Competing Interest Statement

The authors have declared no competing interest.

### Summary of Updates

The revised version of the manuscript accounts for the effects of genetic drift on divergence levels and tipping points, both analytically and via individual-based simulations.

